# The efficacy of resting-state fMRI denoising pipelines for motion correction and behavioural prediction

**DOI:** 10.1101/2024.12.01.626250

**Authors:** Kane Pavlovich, James Pang, Alexander Holmes, Toby Constable, Alex Fornito

## Abstract

Resting-state functional magnetic resonance imaging (rs-fMRI) is a pivotal tool for mapping the functional organization of the brain and its relation to individual differences in behaviour. One challenge for the field is that rs-fMRI signals are contaminated by multiple sources of noise that can contaminate these rs-fMRI signals, affecting the reliability and validity of any derivative phenotypes and attenuating their correlations with behaviour. Here, we investigate the efficacy of different noise mitigation pipelines, including white-matter and cerebrospinal fluid regression, independent component analysis (ICA) – based artefact removal, volume censoring, global signal regression (GSR), and diffuse cluster estimation and regression (DiCER), in simultaneously achieving two objectives: mitigating motion-related artifacts and augmenting brain-behaviour associations. Our analysis, which employed three distinct quality control metrics to evaluate motion influence and a kernel ridge regression for behavioural predictions of 81 different behavioural variables across two independent datasets, revealed that no single pipeline universally excels at achieving both objectives consistently across different cohorts. Pipelines combining ICA-FIX and GSR demonstrate a reasonable trade-off between motion reduction and behavioural prediction performance, but inter-pipeline variations in predictive performance are modest.

## Introduction

Resting-state functional magnetic resonance imaging (rs-fMRI) captures spontaneous fluctuations in the blood-oxygenation-level-dependent (BOLD) signal as individuals remain in a passive state, disengaged from any specific task (Ogawa et al., 1998). Such signals are then used to quantify diverse patterns of inter-regional functional coupling (FC), commonly estimated as correlations between regional signal time courses. rs-fMRI-derived FC has been used extensively to map the intrinsic functional organization of the brain and its disruption in brain disorders (Fox et al., 2005; Smith et al., 2009; Fornito & Bullmore, 2010; Gordon et al., 2017; Kong et al., 2018), as well as how individual differences in various functional network properties relate to behaviour (Hariri, 2009). In turn, FC is frequently associated with measures of cognition, personality, and emotion (Chen et al., 2022; Kong et al., 2019; Ooi et al., 2022; Mueller et al., 2013).

Despite the popularity of rs-fMRI, recent evidence indicates that many correlations between FC and behaviour, as revealed through cross-sectional brain-wide association studies (BWAS), are small and not reproducible in sample sizes typically studied in neuroimaging experiments (i.e., order of hundreds) (Marek et al., 2022). Such small effects can arise under one of two scenarios: (1) the true correlation between brain and behavioural measures is small; or (2) the true correlation is large, but its estimation is attenuated by low reliability and/or validity of the brain variable, the behavioural variable, or both (Tiego et al., 2023; Nikolaidis et al., 2022). The first scenario suggests that cross-sectional correlation analyses are fundamentally limited in their capacity to identify moderate-to-large associations and that we should seek alternative paradigms for relating brain and behaviour. The second scenario suggests that larger associations can be uncovered if we improve the fidelity of the neuroimaging and behavioural measures. Several cost-effective approaches are available for improving the reliability and validity of behavioural measures (for a review, see Tiego et al., 2023). Here, we consider readily applicable methods for improving the fidelity of FC estimates, focusing on the role of different fMRI denoising strategies.

Resting-state fMRI signals are notoriously noisy and susceptible to imaging artifacts, including those induced by head motion, the cardiac cycle, and respiratory variations (Power et al., 2012; Shmueli et al., 2007; Birn et al., 2006). These artifacts reduce the reliability and validity of FC estimates and can attenuate BWAS effect sizes (Teeuw et al., 2021). Many approaches seek to mitigate these effects, with their relative efficacy having been studied extensively (Parkes et al., 2017; Satterthwaite et al., 2013; Ciric et al., 2017; Mascali et al., 2021; Burgess et al., 2016). However, the degree to which these approaches can improve effect sizes in BWAS has received comparatively little attention (for a notable exception focused on one specific processing step, see Li et al., 2019).

To address this question, we explored the effects of six commonly used denoising methods, aggregated into various combinations to define 14 distinct pipelines. We evaluated the performance of each pipeline with respect to measures of both denoising efficacy and the cross-validated prediction of cognitive and personality measures across three independent datasets — the Consortium for Neuropsychiatric Phenomics (CNP; Bilder et al., 2018), the Genomics Superstruct Project (GSP; Holmes et al., 2015, and the Human Connectome Project (HCP; Van Essen et al., 2013). Our comparative approach allowed us to determine whether it is possible to define a single sequence of fMRI processing steps that optimally denoises the data while also maximizing BWAS effect sizes across different psychological measures and datasets.

## Methods

### 1. Participants and data

Resting-state fMRI data in healthy individuals were collected from three publicly available datasets: (i) CNP (*N* = 121) (Poldrack et al., 2016); (ii) GSP (*N* = 1570) (Holmes et al., 2015); and (iii) HCP (*N* = 1200) (Van Essen et al., 2013). The denoising efficacy of different pre-processing pipelines were compared across these datasets of varying imaging quality and parameters. The GSP and HCP datasets were further used to evaluate the efficacy of the pipelines in improving BWAS effect sizes.

The CNP dataset was acquired on two different 3T Trio Siemens scanners at the Ahmanson-Lovelace Brain Mapping Centre and the Staglin Centre for Cognitive Neuroscience. Structural data were acquired using a T1-weighted MPRAGE sequence (TR = 1900 ms, TE = 30 ms, flip angle = 90°) with 176 slices at a 1 mm resolution. Resting-state fMRI data were acquired using a T2*-weighted echo-planar imaging (EPI) sequence (TR = 2000 ms, TE = 30 ms, flip angle = 90°) across 152 volumes with 30 slices. Further details on this dataset can be found in Gorgolewski et al. (2017).

The GSP dataset was acquired on matched 3T Tim Trio scanners at Harvard University and Massachusetts General Hospital. Structural data were acquired using a multi-echo T1-weighted MPRAGE sequence (TR = 2200 ms, TE = 1.5/3.4/5.2/7.0 ms, flip angle = 7°) with 144 slices at a 1.2 mm resolution. Resting-state fMRI data were acquired using a gradient-echo EPI sequence sensitive to BOLD contrast (TR = 3000 ms, TE = 30 ms, flip angle = 85°) across 124 volumes with 47 slices. Behavioural data were collected using a range of different tools including but not limited to the; State-trait anxiety inventory for adults (Speilberger et al., 1970), NEO personality inventory (Costa & McCrae, 2000), and Behavioural inhibition and activation scales (Carver & White, 1994). Details on the specific behavioural items used in this study can be found in Supplementary Table 2. Further details on this dataset can be found in Holmes et al. (2015).

The HCP dataset was acquired on a customized Siemens 3T Skyra at Washington University using a multiband sequence. Structural data were acquired with a T1-weighted MPRAGE (TR = 2400 ms, TE = 2.14 ms, flip angle = 8°). Resting-state fMRI data were acquired using a gradient-echo EPI (TR = 720 ms, TE = 33.1 ms, flip angle = 52°) across 1200 frames. Behavioural data were primarily collected using the National Institutes of Health (NIH) toolbox for assessment of neurobiological and behavioural function (Gershon et al., 2013), as well as supplementary tests that covered fluid intelligence (Penn Matrix Analysis test, Gur et al., 2010), and personality (NEO personality inventory, Costa & McCrae, 2000) among others. Details on the specific behavioural items used in this study can be found in Supplementary Table 3. Further details on the full list of HCP behavioural measures can be found in Van Essen et al. (2012).

### 2. Initial pre-processing

#### 2.1 CNP and GSP initial functional image pre-processing

The CNP and GSP datasets were subjected to a common set of initial pre-processing steps, as implemented in fMRIprep v23.0.2 (Esteban et al., 2019). Briefly, in this standardized pipeline, functional runs were slice-time corrected using Analysis of Functional Neuroimages (AFNI) (Cox, 1996), and realigned to a mean reference image while deriving head motion parameters using FMRIB Software Library (FSL) *mcflirt* (Jenkinson et al., 2002), then distortion correction was performed by co-registering the functional images to the intensity-inverted T1w image using Freesurfer’s *bbregister* (Greve & Fischl, 2009). Images were then resampled to the Montreal Neurological Institute 152 (MNI152) 2009 Non-linear asymmetric template using Advanced Normalisation Tools (ANTs), and the initial four frames from each brain-extracted functional run were excluded to allow for signal stabilisation. Subsequently, each run underwent intensity normalization to a value of 1000, followed by spatial smoothing with a 6 mm Full Width at Half Maximum (FWHM) kernel (with the exception of 24/28P regression pipelines to be described below, where smoothing was conducted after regression). Each T1w image was corrected for intensity non-uniformity with *N4BiasFieldCorrection* (Tustison et al., 2010), skull stripped using FSLs *bet* (Smith et al., 2002), and spatially normalized to the MNI152 2009 Non-linear asymmetric template using a nonlinear registration as implemented with FSLs *fnirt* and *applywarp* (Smith et al., 2004).

#### 2.2 HCP initial functional image pre-processing

The HCP data were initially pre-processed according to the HCP minimal processing pipeline version 3 (Glasser et al., 2013). Specifically, the fMRI volumes underwent a gradient distortion correction before realignment to a single band reference image. Image distortion was corrected using reverse coded spin echo maps and each corrected 3D image was registered to the MNI152 2009 Non-linear asymmetric template using a non-linear transformation obtained using single spline interpolation. The data were then intensity normalized relative to the value of 1000. Further details on the processing steps can be found in Glasser et al. (2013).

#### 2.3 Structural image processing

In all datasets, skull-stripped T1w images in MNI152 space were segmented into white matter (WM), cerebrospinal fluid (CSF), and grey matter (GM) probability maps using the new segment routine from statistical parameter mapping software v8.0 (SPM8). WM and CSF probability maps were then binarized to create tissue-specific masks. As suggested by Power et al. (2017), WM masks were eroded five times and CSF masks twice to avoid any overlap with GM signal. Following erosion, if any mask had less than 5 voxels present the previous erosion cycle was selected as the final mask.

#### 2.4 Functional image processing

In all datasets, six head motion parameters, their temporal derivatives, and the squares of the original and derivative traces were collated (supplied in the HCP dataset and taken from fmriprep in the CNP and GSP datasets) and regressed out to form our baseline pre-processing pipeline. We refer to this step as 24P regression (Parkes et al., 2017). Additionally, these regressors were combined with the averaged WM and CSF signals and their temporal derivatives to form our second baseline pre-processing pipeline (referred to as 28P regression). We then applied distinct denoising procedures to the minimally pre-processed data (Figure 1 & Table 1) before bandpass filtering between 0.008 and 0.08 Hz using a rectangular filter based on the Fourier transform.

**Figure 1.**
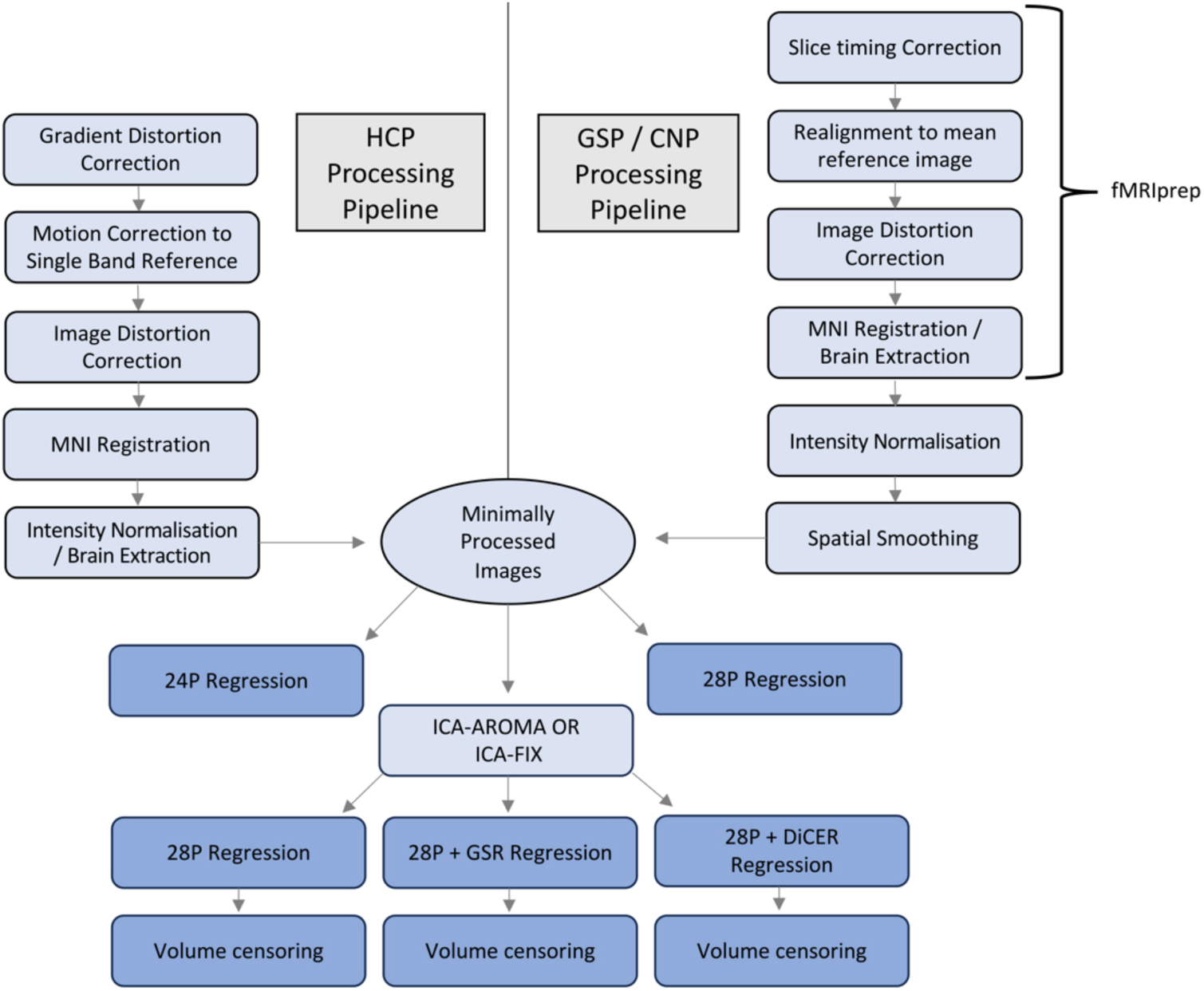
Schematic of the different tools used to form the differential pipelines. The minimal processing pipeline for HCP is shown on the left, while the processing stream for GSP / CNP is shown on the right. Boxes in dark blue indicate final stages that were used in the analysis, while light blue boxes indicate intermediary stages.

**Table 1.**
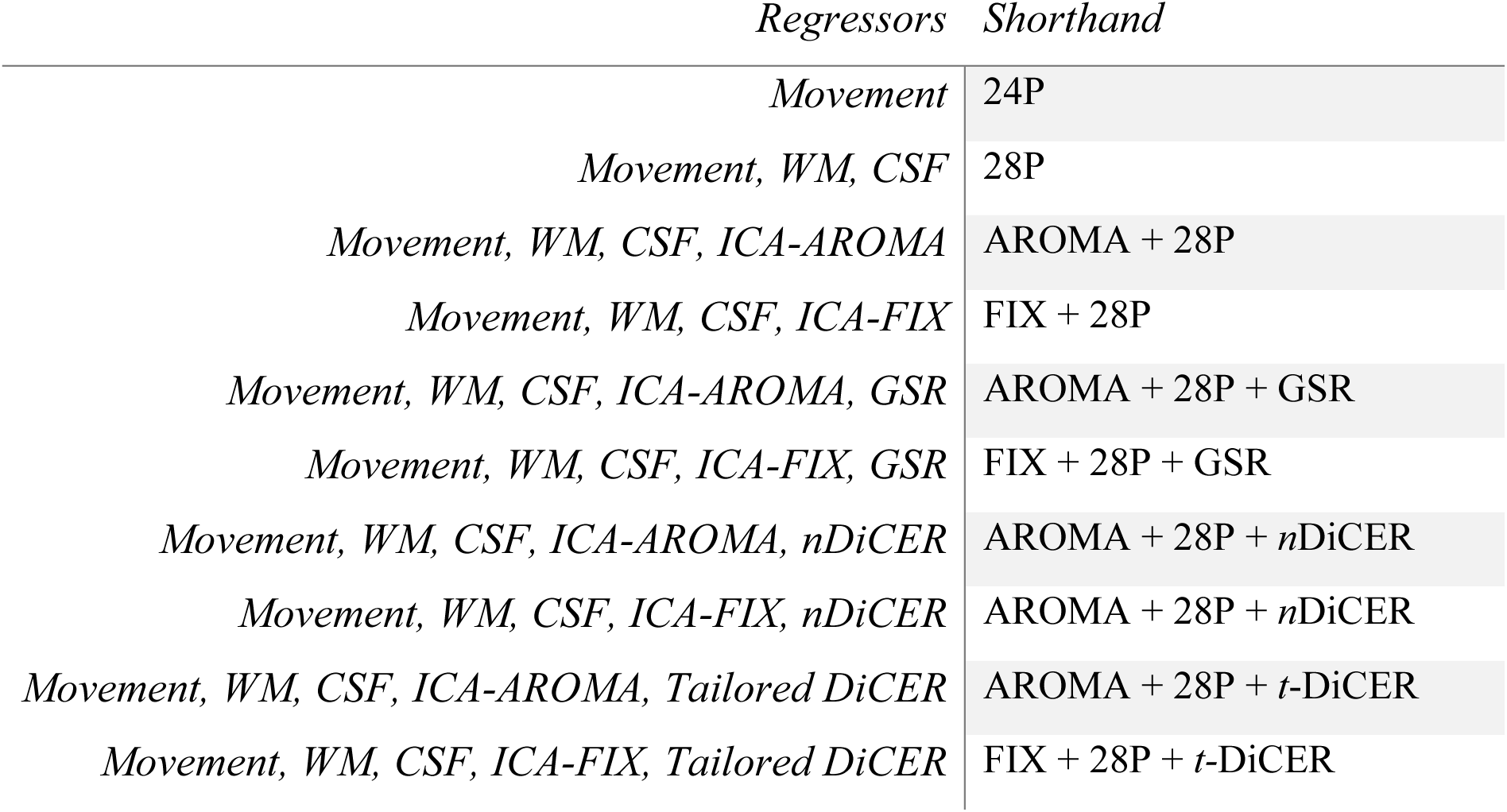
An overview of regressors used and their shorthand. Movement, WM, and CSF regressors were combined with various combinations of ICA-AROMA, ICA-FIX, GSR, or DiCER. Pipelines assessing DiCER were independently evaluated using 1–5 regressors (*n*DiCER), as well as with an individually tailored approach (*t*-DiCER). Each pipeline was also compared with the application of censoring protocols.

| *Regressors* | *Shorthand* |
| --- | --- |
| *Movement* | 24P |
| *Movement, WM, CSF* | 28P |
| *Movement, WM, CSF, ICA-AROMA* | AROMA + 28P |
| *Movement, WM, CSF, ICA-FIX* | FIX + 28P |
| *Movement, WM, CSF, ICA-AROMA, GSR* | AROMA + 28P + GSR |
| *Movement, WM, CSF, ICA-FIX, GSR* | FIX + 28P + GSR |
| *Movement, WM, CSF, ICA-AROMA, nDiCER* | AROMA + 28P + *n*DiCER |
| *Movement, WM, CSF, ICA-FIX, nDiCER* | AROMA + 28P + *n*DiCER |
| *Movement, WM, CSF, ICA-AROMA, Tailored DiCER* | AROMA + 28P + *t*-DiCER |
| *Movement, WM, CSF, ICA-FIX, Tailored DiCER* | FIX + 28P + *t*-DiCER |

### 3. Differential pre-processing pipelines

The minimally pre-processed CNP, GSP, and HCP data were then subjected to different de-noising pipelines. We first considered two different denoising approaches:

1. Independent Component Analysis based strategy for Automatic Removal of Motion Artifacts (ICA-AROMA) denoising (Prium et al., 2015); and
2. ICA-based X-noiseifier (ICA-FIX) denoising (Salimi-Khorshidi et al., 2014).

We focused on these two approaches because head-to-head comparisons have indicated that ICA-AROMA does well on single-band data (Parkes et al., 2017; Aquino et al., 2020; Ciric et al., 2017), while ICA-FIX represents an extension of the approach that should better parse signal and noise and is the standard denoising algorithm applied to the widely used HCP dataset.

Each of the following three approaches was then run to address wide-spread signal deflections (WSDs) (sometimes also called global signal fluctuations):

1. No WSD removal;
2. Global signal regression (GSR) using the mean signal estimated from the whole brain mask; and
3. Removal of WSDs via diffuse cluster estimation and regression (DiCER) (Aquino et al., 2020).

Hence, we apply six possible combinations of denoising and WSD removal algorithms along with our two baseline pre-processing strategies. We further ran each with and without censoring of high motion frames, yielding a total of 14 distinct pipelines (Supplementary Table 1). In the following, we explain the details of how each of these procedures were implemented.

#### 3.1 ICA-AROMA

ICA-AROMA (Prium et al., 2015) relies on spatial ICA to automatically remove components largely related to motion. The approach uses a classifier to label components as noise for subsequent removal. The criteria for identifying noise components is based on four features: the amount of high-frequency content in the component time series; the correlation between component and motion-related time courses; the amount of activation in the spatial features of the component that is attributable to brain edges; and the level of component activity contained in CSF.

We applied ICA-AROMA to both the CNP and GSP datasets, Given that ICA-AROMA was trained using single-band data, it was not used on the HCP dataset, which employs multiband sequences. We used the non-aggressive variant, which only removes the variance unique to noise components that are not shared with any of the signal components (Prium et al., 2015). As ICA-AROMA is non-deterministic, the components and resulting FC estimates can change between runs (Supplementary Figure 1). We therefore ran the algorithm 10 times to assess algorithmic variability.

#### 3.2 ICA-FIX

In contrast to ICA-AROMA, ICA-FIX requires user-derived classifications of signal and noise components to train a classifier in a subset of the data. Despite being labour-intensive, this process allows for more precise removal of noise components while preserving as much neural signal as possible, compared to the fully automated approach of ICA-AROMA (Salimi-Khorshidi et al., 2014; Griffanti et al., 2014). ICA-FIX was applied to both the HCP and GSP datasets, as the CNP dataset had a low sample size and it wasn’t feasible to split into train and test sets for further analysis. The HCP data had undergone prior ICA-FIX processing (for additional details, refer to Glasser et al., 2013). We implemented ICA-FIX on the GSP data using a similar approach. We first established a held-out training set of 40 subjects (20 males; mean age 22.05 years) and ran spatial ICA separately in each individual using FSL’s melodic (Beckmann & Smith, 2004). We then manually classified the resulting components as either signal or noise. ICA-FIX assigns a probability value indicating the likelihood that a component is noise. To determine an optimal cut-off threshold for this probability, we conducted leave-one-out testing on the training set across various thresholds and evaluated the classifier’s performance (Supplementary Figure 2). Based on these evaluations, we selected a threshold of 40, which yielded a mean True Positive Rate of 91.7% and a mean True Negative Rate of 89%.

#### 3.3 Global Signal Regression

GSR was executed by regressing the average time series across the entire brain from each voxel in the Blood-Oxygen-Level-Dependent (BOLD) data. Following ICA-based denoising, whole-brain masks were applied to derive the mean global signal across all voxels. Subsequently, the mean global signal and its first derivative were regressed out with the WM and CSF signals and their respective derivatives.

#### 3.4 DiCER

DiCER is a data-driven method designed to remove WSDs from voxel-wise BOLD time series by identifying and regressing out clusters of weakly correlated voxels (Aquino et al., 2020). DiCER employs a density-based spatial clustering algorithm to identify highly correlated/anticorrelated clusters of voxels. Once clusters are identified, DiCER calculates an adjusted mean signal for each cluster by determining a central voxel for the cluster and flipping the signs of time series for voxels that are anticorrelated with this central voxel. The estimated WSD is then removed from each voxel’s time series via linear regression. This process, based on the original implementation by Aquino et al. (2020), is repeated until no new clusters are found or a maximum of five iterations is reached. The approach thus removes WSDs without the use of GSR, which has been shown to introduce artificial anti-correlations and bias FC estimates (Fox et al., 2009; Saad et al., 2012). Relative to GSR, DiCER is comparable or more effective at denoising, while preserving variance likely to be of neuronal origin (Aquino et al., 2020).

A limitation of DiCER is that the optimal number of iterations remains unknown. We therefore assessed the impact of running DiCER with between 1 and 5 iterations, which we will refer to as *n-*DiCER, where *n* refers to the number of iterations used. We also evaluated a heuristic strategy for tailoring the number of iterations for each individual by selecting the number of iterations that maximizes the modularity of each individual’s resulting FC network, which we called tailored DiCER (t-DiCER; see Supplementary Information for further details). This strategy follows evidence that brain networks tend to favour a modular organization (Sporns & Betzel, 2016).

### 4. Volume censoring

Researchers will often use censoring procedures to identify fMRI volumes that exceed a certain motion threshold (Siegel et al., 2014). While it is seen as an effective method for removing motion-contaminated frames, it comes at the expense of removing large amounts of data.

In our analyses, we censored specific high-motion frames based on their level of framewise displacement (FD). FD is a metric that summarizes frame-to-frame changes in head motion based on six motion parameters describing translations and rotations (Power et al., 2014). In the GSP and CNP datasets, FD was calculated through fmriprep on the unprocessed functional scans of individual participants, following the method described by Power et al. (2014). This approach was similarly applied to calculate FD in the HCP dataset, where additionally, FD traces were bandpass filtered between 0.31 Hz and 0.43 Hz following the guidelines of Fair et al. (2020). This filtering was done to mitigate the impact of respiratory artifacts on motion estimates, which are typically influenced by multiband sequences.

Frames exceeding an FD threshold of 0.3 mm were flagged for censoring. Additionally, if there were fewer than five consecutive frames between any initially flagged frames, those frames were also marked for censoring. Subsequently, interpolation of censored frames was performed using the Lomb-Scargle Periodogram method, following the approach detailed by Power et al. (2014). Censoring was performed after confound regression but before bandpass filtering.

### 5. FC estimation

Pre-processed fMRI data were multiplied by the corresponding grey matter probability map for each participant (see Section 2.3) so that a mean time series, weighted by each voxel’s grey matter-probability, was extracted for each of 300 cortical regions, as defined using a widely used parcellation (Schaefer et al., 2018). We then calculated the Pearson’s correlation between the time courses of each pair of regions and normalized the correlation distribution using Fishers *r*-to-*z* transformations.

### 6. Subject exclusion

Participants with high levels of motion, as quantified using FD, were excluded from analyses. Following the recommendations of Satterthwaite et al. (2013) and Parkes et al. (2017), participants were excluded if any of the following criteria were met: (i) mean FD > 0.3 mm; (ii) more than 20% of FDs were above 0.2 mm; and (iii) any FD > 5 mm. Participants were also excluded if more than half of their frames were censored.

### 7. Quantifying denoising efficacy

We quantified the denoising efficacy of each pipeline using three quality control (QC) metrics, as per Parkes et al. (2017): QC-FC correlations; QC-FC distance-dependence; and the variance explained by the first principal component of the FC matrix (VE1).

QC-FC correlations correspond to the cross-participant correlation between each individual’s mean FD and FC estimates at each entry, or edge, of the FC matrix. We thus obtained one QC-FC correlation for each of 44,850 edges between each pair of 300 regions. This correlation quantifies the residual relationship between FC and head motion.

QC-FC distance dependence examines how QC-FC correlations vary as a function of the physical distance between each pair of regions, given prior evidence that in-scanner movement spuriously increases short-range coupling relative to medium and long-range coupling (Van Dijk et al., 2012). We quantified this dependence as the Spearman rank correlation between the QC-FC correlation of each pair of regions and the Euclidean distance between the corresponding region centroids (Power et al., 2015).

VE1 was estimated using principal components analysis (PCA) of each individual’s variance-covariance matrix. PCA decomposes the matrix into a set of linearly orthogonal components, ordered by the variance that each explains. Data dominated by WSDs, which often (although not always) arise from head motion and respiratory variations (Aquino et al., 2020; Power et al., 2014), will show high levels of global synchrony across the brain, meaning that most of the variance will be explained by the first component. Successful removal of WSDs will thus reduce VE1 estimates (Aquino et al., 2020).

VE1 was estimated by dividing the first eigenvalue of the PCA by the sum of all eigenvalues. Note that lower VE1 estimates are not always better. A very aggressive denoising procedure will remove any structure in the data, and the FC matrix will resemble white noise. In this case, VE1 will be very low. The optimal value of VE1 for brain networks is unclear, but the quantity offers a useful indication of the degree to which a given denoising procedure has removed globally coherent WSDs in the non-denoised data (Aquino et al., 2020).

### 8. Behavioural measures

Both the GSP and HCP datasets provide a range of behavioural measures that span cognition, emotion, and personality. Following the analysis conducted in Li et al. (2019), we selected the same 22 behavioural measures from the GSP dataset and 58 from the HCP they used (see Li et al. (2019) for details). Participants who had any of these behaviours missing were removed from further analysis, such that only participants who had all behaviours analysed in their dataset were included (Figure 2). Prior to prediction, we regressed age, sex, and FD from each behavioural measure. The regression coefficients were calculated from the training set and then applied to the test set to avoid leakage between train and test sets in the prediction model (see below) (Chyzhyk et al., 2022).

**Figure 2.**
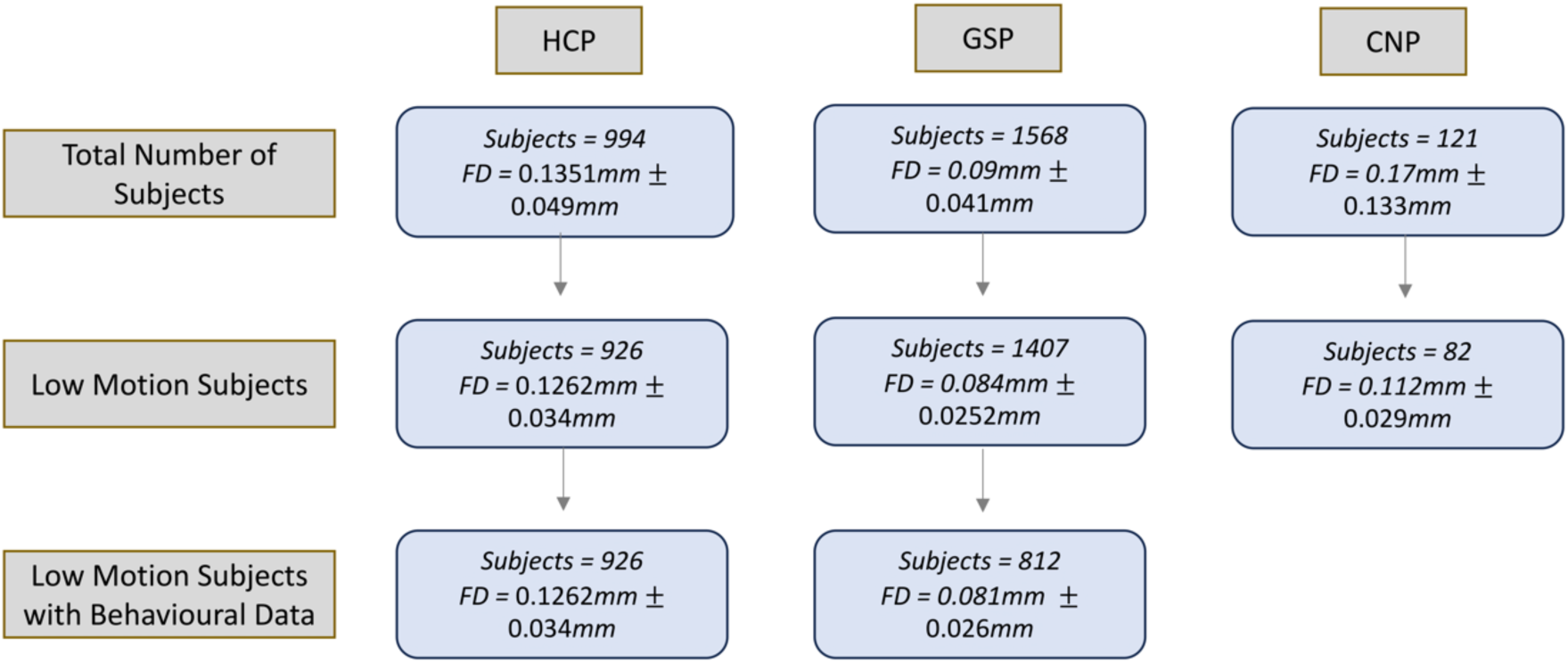
Participant exclusions based on FD and censoring exclusions across the HCP, GSP, and CNP datasets. The censoring exclusion criteria were: mean FD < 0.3 mm; more than 20% of FDs higher than 0.2 mm; any FD > 5mm; or more than half of frames censored. The mean and standard deviation of the FDs across participants are shown.

#### 8.1 Behavioural prediction

We assessed the ability of FC estimates derived from each pipeline to predict behaviour using kernel ridge regression (KRR) with a 20-fold cross-validation, following the implementation outlined in Li et al. (2019). KRR was chosen to facilitate comparison with Li et al. (2019) and because of its robust performance in brain-behaviour predictions (He et al., 2020). FC estimates were used to predict 23 raw behavioural scores, encompassing psychometric and cognitive measures, from the GSP dataset and 58 scores covering cognition and emotion from the HCP dataset. HCP participants from the same family were included in the same fold to avoid train-test predictions across members from the same family.

KRR requires selection of an L2 regularization parameter, which adds a penalty equivalent to the sum of the squared values of the weights to the loss function. By doing so, it controls the trade-off between achieving a low training error and a low validation error, preventing overfitting. We selected the value of this parameter using an additional 20-fold nested cross-validation on each of the initial 20-folds, such that for each fold an additional 20-fold cross validation was applied to find the optimal value for the regularisation parameter. Each behavioural prediction was repeated 20 times, and the average accuracy was taken across repetitions to ensure the stability of our results (Kong et al., 2019).

### 9. Data and Code Availability

Neuroimaging and behavioural data from all datasets are publicly available and can be downloaded from the following links: CNP (https://openneuro.org/datasets/ds000030/versions/00016); GSP (https://www.neuroinfo.org/gsp); and HCP (https://www.humanconnectome.org/study/hcp-young-adult/document/1200-subjects-data-release).

All codes used in our analysis, including links and references to code employed from others, can be found at https://github.com/kanepav0002/rsfmri_prediction

## Results

### 1. Data Characteristics

We first excluded high motion subjects based on a set of stringent FD and censoring thresholds (Figure 2). Mean FD was highest in the HCP dataset, followed by CNP, and GSP was last. Note that the HCP data were acquired with a multiband sequence, which has been shown to inflate FD traces relative to single-band acquisitions (Fair et al., 2020; Power et al., 2019). While we filtered out respiratory frequencies to mitigate some of this inflation, caution should be taken when comparing FD traces across single and multiband data acquisitions. For the behavioural analysis, subjects were only included if they had no missing data for the 22 behaviours analysed in the GSP dataset and the 58 in the HCP dataset (Figure 2).

### 2. QC-FC correlations

We first examined the denoising efficacy of each pipeline using QC-FC correlations across the CNP, GSP, and HCP datasets. A successful de-noising pipeline should exhibit a QC-FC distribution centred around zero with little dispersion.

Figure 3 shows that, across datasets, ICA-AROMA and ICA-FIX reduced the magnitude of QC-FC correlations relative to pipelines relying only on either 24P or 28P regression. However, the biggest reductions occurred when these pipelines were combined with some form of WSD removal; i.e., either GSR or DiCER. One iteration of DiCER shifted the mode of the QC-FC distribution into the negative range, whereas two or more iterations yielded an approximately zero-centred distribution, similar to the effect of GSR. Increasing the number of iterations to five slightly truncated the distribution tails, suggesting greater denoising efficacy with a more aggressive application of the technique. The performance of t- DiCER was somewhere in between 1-DiCER and 5-DiCER (for the number of iterations selected for each person, see Supplementary Figure 3). Censoring of high-motion frames did not have an appreciable impact on QC-FC distributions.

**Figure 3.**
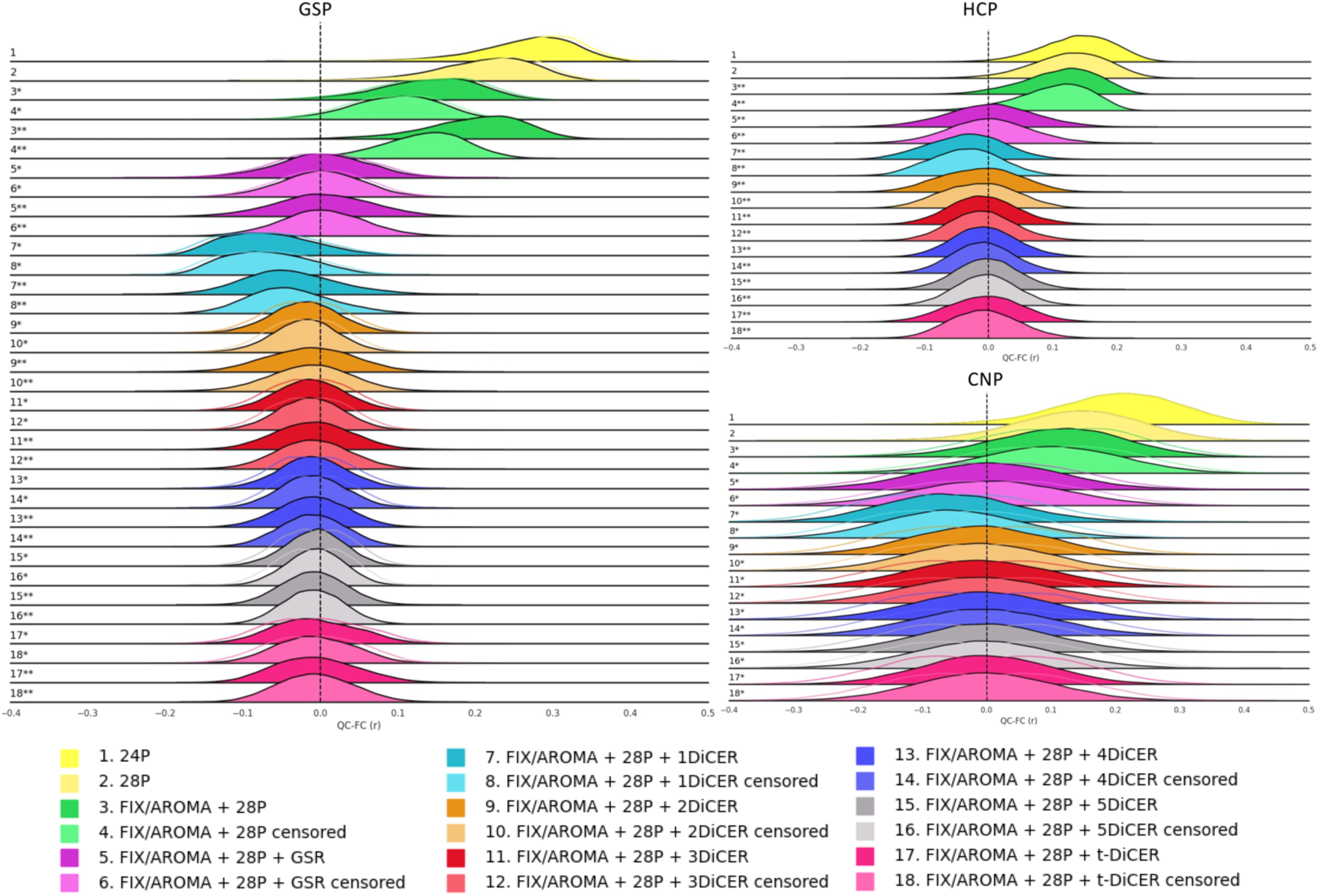
Distribution of QC-FC values (Pearson’s *r*) across all edges for each pipeline and dataset employed, averaged across participants. For pipelines utilizing ICA-AROMA, the mean QC-FC result is depicted across 10 different runs, with non-shaded distributions representing the minimum and maximum QC-FC results across runs (also shown in Supplementary Figure 4). Pipelines incorporating ICA-AROMA are labelled with a single asterisk, while those using ICA-FIX are labelled with a double asterisk.

We did not observe major differences between ICA-AROMA and ICA-FIX in the GSP dataset, which was the only dataset subjected to both algorithms. Additionally, the algorithmic variability of ICA-AROMA was low when combined with 28P regression, but substantially increased with a higher number of DiCER iterations (Supplementary Figure 4). For instance, in the CNP dataset with 28P alone, the mean QC-FC ranged between 0.073 and 0.145, but following 1-DiCER it ranged between −0.105 and −0.007 and between −0.069 and 0.064 following 5-DiCER.

### 3. QC-FC distance dependence

Figure 4 shows that the efficacy of each pipeline in mitigating the distance dependence of QC-FC correlations was variable across datasets. In the CNP dataset, ICA-AROMA with or without GSR or DiCER generally reduced the distance-dependence of QC-FC estimates relative to the simple 28P regression. However, some specific implementations of DiCER (specifically 3-DiCER and t-DiCER) were associated with slight increases in distance-dependent correlations. The inclusion of motion censoring was also associated with a slight increase in distance-dependence for either t-DiCER or DiCER with more than two iterations. The results obtained in the HCP dataset were similar, with distance-dependent QC-FC increasing with more iterations of DiCER. In the GSP dataset, ICA-AROMA increased QC-FC distance-dependence compared to ICA-FIX. GSR and DiCER were also associated with increased distance-dependence compared to pipelines that did not incorporate these steps. The only exceptions were ICA-AROMA + 28P + 2DiCER + censoring and ICA-FIX + 28P + 3DiCER.

**Figure 4.**
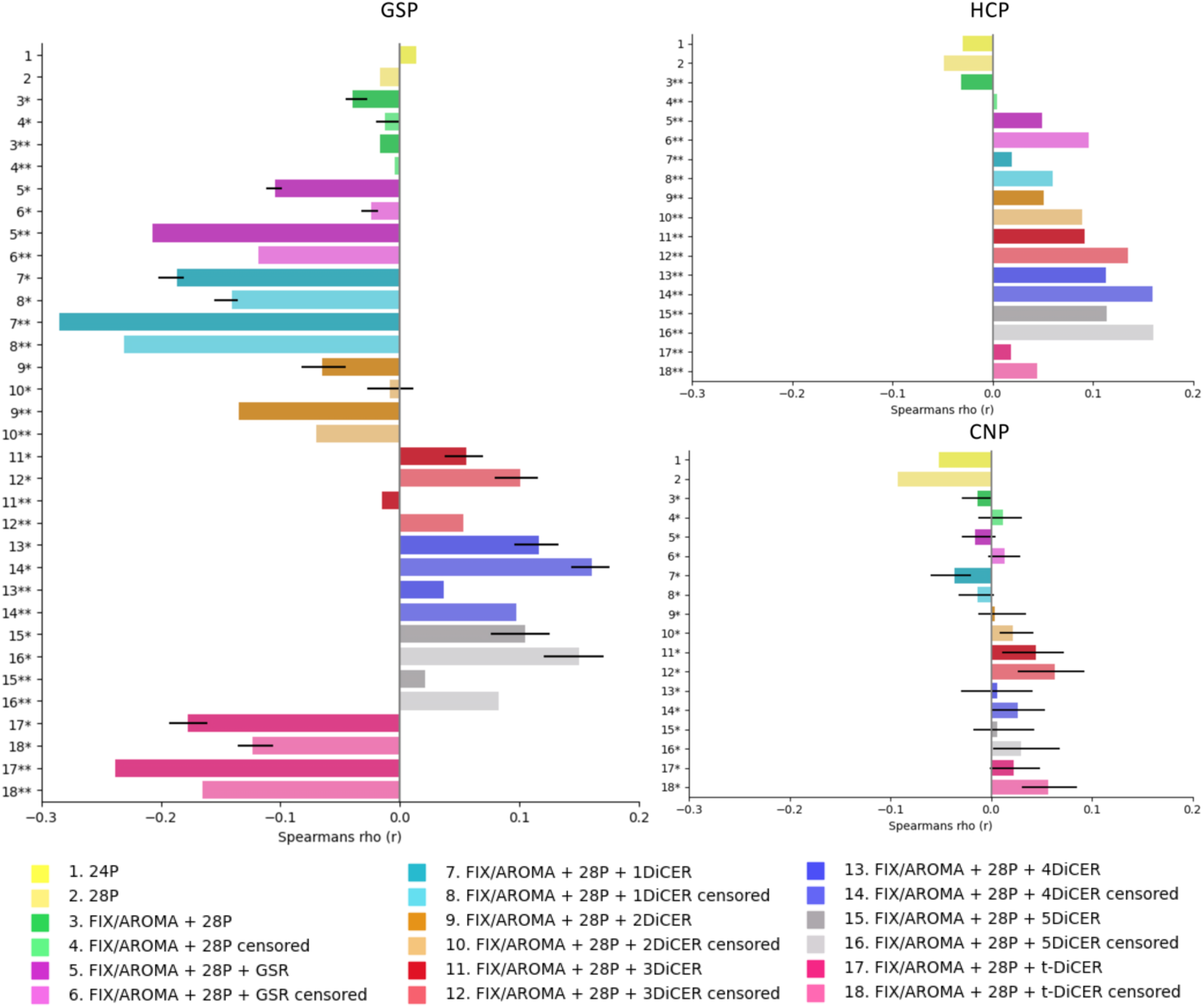
QC-FC distance-dependent correlations for each pipeline and dataset, averaged across participants. For pipelines using ICA-AROMA, bars correspond to the mean across runs and error bars represent the minimum and maximum range across runs. Pipelines incorporating ICA-AROMA are labelled with a single asterisk, while those using ICA-FIX are labelled with a double asterisk.

The algorithmic variability of ICA-AROMA was comparable to the variations observed for QC-FC correlations, and increased when either GSR or DiCER were included in the pipeline (see error bars in Figure 4). This trend was more salient in the CNP dataset compared to the GSP dataset.

### 3. Variance explained by PC1

VE1 can be used to quantify the degree to which each pipeline has removed globally coherent structure driven by WSDs in the FC matrix (Figure 5). As with QC-FC, VE1 was attenuated by the inclusion of either ICA-AROMA or ICA-FIX across all datasets. This reduction was more dramatic following further inclusion of either GSR or DiCER. Increasing the number of DiCER iterations progressively reduced VE1, which is expected given that the algorithm iteratively identifies and removes large-scale structures in the data. t- DiCER yielded VE1 estimates in between those obtained with one and five iterations. Censoring led to a greater attenuation of VE1 in pipelines with solely ICA + 28P denoising methods, but this variation was limited when either GSR or DiCER was applied. Differences between ICA-AROMA and ICA-FIX were minimal, and inclusion of censoring did not have a major impact on VE1.

**Figure 5.**
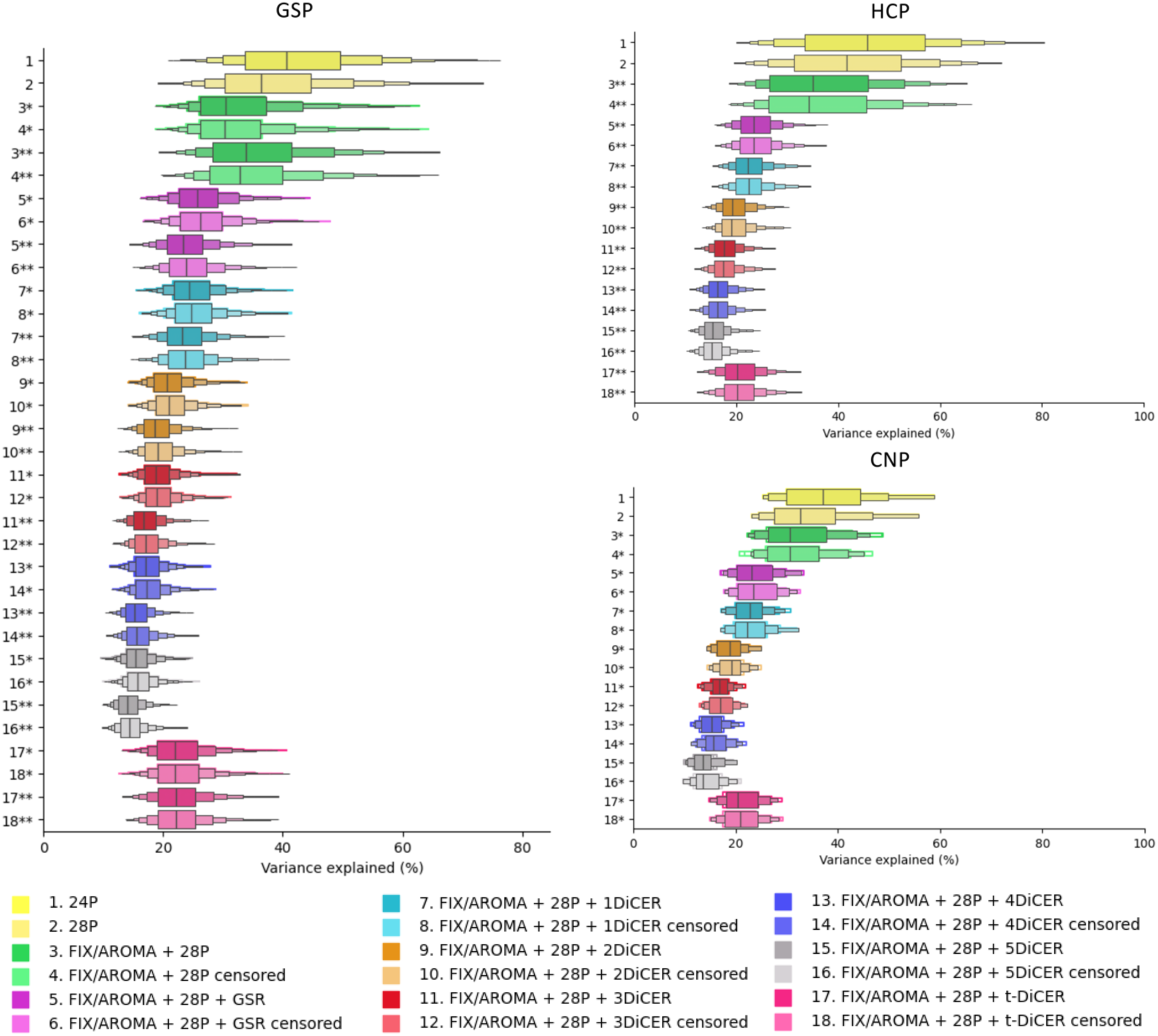
Distributions of VE1 estimates for each pipeline and dataset. The middle of each box represent the 50% percentile and the outer boxes represent decreasing percentiles (i.e., 25%, 12.5%, and 6.25%). For pipelines using ICA-AROMA, the mean minimum and maximum result is shaded around each mean result. Pipelines incorporating ICA-AROMA are labelled with a single asterisk, while those using ICA-FIX are labelled with a double asterisk.

### 4. Behavioural prediction

Measures of denoising efficacy, such as QC-FC-based metrics and VE1, can be used to quantify the degree to which a pipeline has successful removed noise in the data. However, there is no guarantee that pipelines performing well on such metrics yield more reliable or valid FC estimates since a very aggressive denoising procedure will generally minimize denoising efficacy metrics while also potentially removing signals of interest. It remains unclear how the preservation of such signals can be best quantified and several different approaches have been considered (Glasser et al., 2018; Aquino et al., 2020; Li et al., 2019). Here, following Li et al. (2019). Here, we evaluated the ability of FC estimates obtained under each denoising pipeline to predict different measures of behaviour acquired outside the scanner, directly assessing the degree to which each denoising approach influences BWAS effect sizes. Specifically, we evaluated each pipeline’s efficacy in maximizing the variance explained in behaviour using multivariate Kernel Ridge Regression (KRR) in the HCP (58 behavioural measures) and GSP (22 behavioural measures) datasets.

Figure 6 shows the mean and interquartile ranges averaged across all KRR repetitions, cross-validation folds, and measures, or after splitting the measures into those of cognition or personality (details of each behaviour variable are found in Supplementary Tables 2 and 3). In the HCP dataset, cognitive measures were most strongly predicted by ICA-FIX + 28P + GSR without censoring followed by ICA-FIX + 28P + 1DiCER with censoring (mean *r* = 0.17 and *r* = 0.167, respectively). Increasing the number of DiCER iterations generally reduced model performance and censoring had little impact relative to the uncensored pipelines. The most aggressive denoising pipeline, i.e., ICA-FIX-28P + 5DiCER, was associated with the lowest prediction accuracy. For personality measures, ICA-FIX + 28P + GSR (mean *r* = 0.077) was the best performing pipeline, followed by the 2DiCER pipeline (mean *r* = 0.068). Effect sizes for personality measures were generally small, with *r* = 0.054 on average.

**Figure 6.**
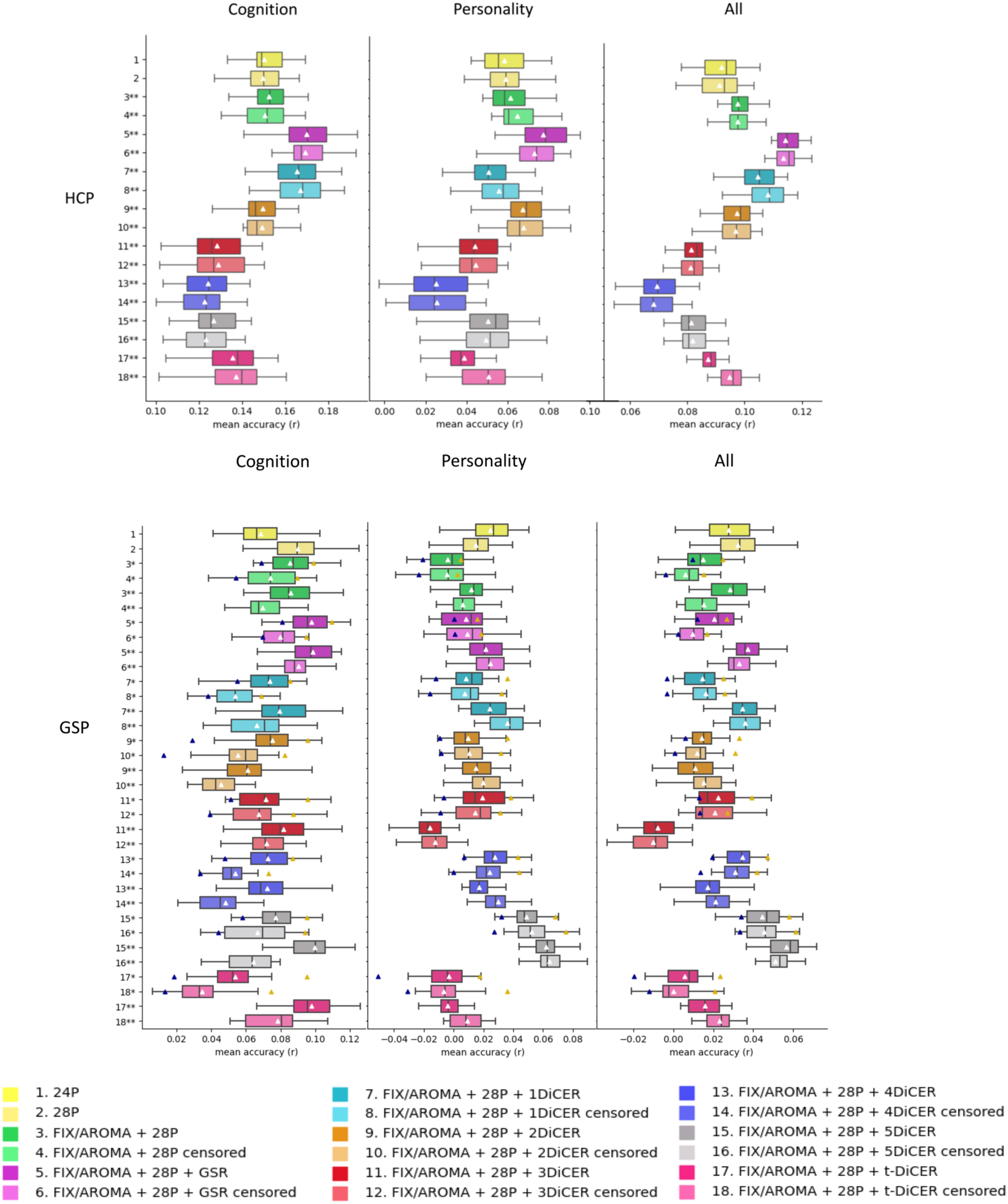
Mean Kernel Ridge Regression (KRR) prediction accuracies averaged across cognitive, personality, and all behavioural measures in the GSP and HCP datasets. Boxes depict the medians and interquartile ranges across 20 cross-validation splits and white triangles show the mean. For pipelines incorporating ICA-AROMA, the result shows an average across 10 different runs. Mean minimum performance across runs is represented by a blue triangle and mean maximum performance by a yellow triangle. Pipelines incorporating ICA-AROMA are labelled with a single asterisk, while those using ICA-FIX are labelled with a double asterisk.

Considering cognitive measures in the GSP dataset, ICA-FIX + 28P + 5-DiCER was associated with the largest effect size, but the increment in performance was marginal to that obtained with 28P regression alone, increasing to a mean *r* = 0.099 from a mean *r* = 0.069. ICA-FIX or ICA-AROMA with 28P and GSR, and ICA-FIX with t-DiCER, performed similarly. DiCER with 2, 3, or 4 iterations generally showed poorer performance than pipelines with GSR. Censoring also reduced prediction performance.

As with the cognitive measures, ICA-FIX + 28P + 5-DiCER yielded the largest effect sizes for predicting personality scores in the GSP dataset, although all effect sizes were smaller, being *r* = 0.017 on average and never exceeding *r* = 0.064. Prediction results varied considerably across different runs of ICA-AROMA, especially when combined with multiple iterations of DiCER (as outlined by the blue and yellow triangles in Figure 6). For instance, mean effect sizes fluctuated between *r* = 0.058 and *r* = 0.095 when using 5-DiCER to predict the cognitive variables, suggesting that stochastic fluctuations in the implementation of the algorithm should be accounted for in BWAS.

## Discussion

Here we assessed the efficacy of six different fMRI denoising procedures (WM/CSF regression, ICA-AROMA, ICA-FIX, GSR, DiCER, and censoring) aggregated into 14 distinct pipelines in mitigating motion- and WSD-related contamination of FC estimates across three independent datasets. We also explored the impact of each pipeline on BWAS effect sizes in two of the datasets. Our findings indicate that (i) there is no single pipeline that optimally denoises all datasets while maximising BWAS effect sizes; (ii) variations in the impact of denoising on BWAS effect sizes are minor; (iii) GSR notably decreases motion confounds and VE1, and often leads to a slight improvement of behavioural predictions when combined with ICA-FIX; and (iv) DiCER effectively reduces motion-related confounds and VE1, but the optimum number of iterations for maximising BWAS effect sizes is dataset-dependent. In the following, we consider how our findings can inform choices around the inclusion (or not) of different processing procedures in study-specific pipelines.

### ICA-based de-noising

ICA-based methods are among the most popular strategies for mitigating motion related information in fMRI data, either in the form of ICA-AROMA or ICA-FIX. Our findings reveal that neither method offers a major reduction of QC-FC correlations or VE1 compared to 28P regression alone, although they do reduce QC-FC distance-dependence to some extent. ICA-based denoising on its own may thus be insufficient to remove motion-related and respiration-related contributions to resting-state FC estimates, as suggested by past work (Parkes et al., 2017; Aquino et al., 2020; Ciric et al., 2017, Power et al., 2015).

It is informative to consider these results in relation to behavioural predictions, where we found that the accuracies of ICA-FIX + 28P and ICA-AROMA + 28P pipelines were slightly less than 28P regression alone in the GSP dataset. This result suggests that ICA-based denoising may remove some behaviourally relevant variance from FC estimates, although this effect is small.

Overall, all pipelines that incorporated ICA-FIX offered comparable denoising performance to ICA-AROMA across QC-FC metrics, along with a slight improvement in VE1. In terms of predicting behaviour, pipelines incorporating ICA-FIX almost always outperformed ICA-AROMA. Since ICA-FIX is trained specifically on the dataset being used (Salimi-Khorshidi et al., 2014; Griffanti et al., 2014), it is better able to remove noise that is specific to the dataset, as evidenced by the reduction in VE1. The manual training also affords enhanced sensitivity to identifying a wider range of noise-related components than the heuristic classifications used by ICA-AROMA. The increased denoising efficacy of ICA-FIX may then drive the slight increase in effect sizes that we observed in behavioural predictions.

An important consideration when using ICA-AROMA is algorithmic variability. The non-deterministic nature of the spatial ICA decomposition means that the resulting FC estimates can vary from run to run. We found that performance across a range of quality control metrics can vary considerably across different runs of the algorithm. This variability is particularly pronounced in behavioural predictions, where the mean performance showed ∼two-fold variation across iterations. Future work using ICA-AROMA should thus run the algorithm multiple times and report distributions for effect size estimates. Note that ICA-FIX also depends on spatial ICA and is thus also likely to show some degree of algorithmic variability. We did not consider this effect here due to the higher computational burden of ICA-FIX, but future work may seek to explore the degree of this variability.

### Global Signal Regression (GSR)

The use of GSR remains a contentious issue in fMRI research. On the one hand, GSR has demonstrated efficacy in reducing motion-related artifacts and improving brain-behaviour associations (Power et al., 2014; Li et al., 2019). On the other hand, it introduces spurious negative correlations and has the potential to eliminate neural information relevant to the signal of interest (Fox et al., 2009), although some studies suggest that it actually preserves neural information (Matsui et al., 2016; Li et al., 2019). Our results indicate that employing GSR following ICA-based denoising reduces QC-FC and VE1, but its effects on QC-FC distance-dependent correlations were contingent on the dataset, leading to overall reductions in the HCP and CNP datasets while increasing short-range QC-FC in the GSP dataset. When combined with ICA-FIX + 28P, GSR improved BWAS effect sizes compared to 28P alone.

These findings corroborate prior research highlighting GSR’s ability to attenuate artifacts stemming from non-neuronal physiology and to strengthen associations between FC and behavioural measures (Li et al., 2019; Power et al., 2014; Byrge & Kennedy, 2018). Studies in mice also indicate that GSR increases the correspondence between BOLD and Ca+ dynamics, suggesting that it removes noise while preserving neural activity (Matsui et al., 2016). Recent work has further shown that a large fraction of the global signal can be attributed to breathing and cardiac-related fluctuations, while only a small amount can be attributed to electrophysiological brain activations (Xifra-Porxas et al., 2024). These considerations suggest that GSR is an effective de-noising tool that can improve, albeit marginally, BWAS effect sizes.

### Diffuse Cluster Estimation and Regression (DiCER)

DiCER iteratively identifies and regresses out weakly correlated groups of grey matter voxels across fMRI time series, offering a data-driven alternative to GSR (Aquino et al., 2020). The approach thus aims to provide similar benefits to GSR without necessarily imposing artificial structure on the data (Fox et al., 2009). Our findings reveal that DiCER with one, two, or a tailored number of iterations achieve comparable or superior de-noising performance to GSR in terms of removing motion-related artifacts (as indicated by lower QC-FC correlations) and reducing VE1. Performance on these metrics generally improves with more iterations, as is expected given that each iteration progressively removes more WSDs from the data.

The effects of DiCER on QC-FC distance-dependence were contingent on the dataset. In the GSP dataset, all iterations of DiCER increased this distance-dependence relative to the baseline processing of 24P. In the HCP and CNP datasets, one to two iterations of DiCER reduced the distance-dependence, but three to five iterations increased it, relative to the baseline processing of 24P.

Despite this denoising efficacy, no DiCER variant ever surpassed GSR in enhancing mean BWAS effect sizes in the HCP dataset, even when using 1-DiCER, which shares many conceptual similarities with GSR (Aquino et al., 2020). This result suggests that GSR may better preserve behaviourally relevant variance in FC estimates, although in many cases the difference is marginal. Given the equivalent or slightly superior denoising efficacy of DiCER, these findings suggest that the choice between the two approaches may depend on how one trades off these two factors. Our results suggest that this choice must be made on a case-by-case basis for each dataset.

The use of modularity maximization as a heuristic for tailoring the number of DiCER iterations to individual participants showed good denoising efficacy but near-poorest BWAS effect sizes. This is perhaps surprising given evidence that brain networks, like other complex systems, exhibit a preference for highly modular organization that is believed to facilitate specialized processing, support complex neural dynamics, and minimize network wiring costs (He et al., 2009; Betzel et al., 2017). Prior research has also indicated that the modularity of a brain network can serve as a predictor of learning and cognitive performance (Weinberger et al., 2022; Baniqued et al., 2019). Despite these desirable properties, modularity maximization alone appears to be insufficient for optimizing DiCER-based denoising pipelines. This may be due to the fact that modularity maximization is a relatively simple heuristic that may be inferior to alternative approaches that use exact methods compared to heuristic methods to partition complex networks into coherent modules (Aref & Mostajabdaveh, 2024). These algorithms may provide more effective in individualizing the DiCER algorithm.

### Optimal pipelines across datasets

One notable finding in this analysis is that the optimal pipelines for enhancing behavioral predictions differed between datasets. Specifically, ICA-FIX + GSR was optimal for the HCP dataset, while ICA-FIX + 5-DiCER performed best for the GSP dataset. Differences in the level of noise of each dataset may play a role. The GSP dataset may contain higher levels of either motion-related or non-motion related noise than the HCP data. Although comparisons of mean FD and VE1, which are sensitive to both sources of noise, in the minimally processed data suggest that the opposite is the case, comparisons are complicated by differences in scanner strengths, sampling rates, and acquisition lengths of the data (Power et al., 2019; Fair et al., 2020). Indeed, it is well-established that scan times exceeding 10 minutes are necessary to produce stable and reproducible within-subject FC matrices (Gonzalez-Castillo et al., 2014). The HCP dataset has a scan time of 20 minutes, while the GSP dataset used in our final analysis has participants with 6 (207 participants) and 12 minutes (605 participants) of resting-state data. As the large part of this sample contains data under the 10 minute threshold, FC estimates within the GSP dataset may not be as stable as those for the HCP data. Notably, it has recently been shown that prediction performance increases as scan duration goes up, and that this effect can continue beyond 20 minutes of scan time (Ooi et al., 2024). This low scan duration may account for the lower overall effect sizes and increased need of further denoising in the GSP sample, and highlights the need for long scan times in BWAS studies.

### Does denoising improve BWAS effect sizes

One motivation for our analysis was to investigate whether optimizing fMRI denoising pipelines can improve BWAS effect sizes beyond the low baseline (mean effect of *r* = 0.17) reported in prior work (Marek et al., 2022). Our findings indicate that while some pipelines can increase effect sizes, the gains are small and the optimum pipeline for one dataset or behavioural measure may not be optimal for a different dataset. As such, fMRI denoising, on its own, may be insufficient to significantly improve BWAS effect sizes and, if taken in isolation, our findings cannot definitively determine whether small BWAS effect sizes are due to a small true correlation or reflect attenuation effects caused by the poor reliability and/or validity of our measures (Tiego et al., 2023; Nikolaidis et al., 2022). A combination of different strategies to improve data fidelity (DeYoung et al., 2024), which includes refining the precision of behavioural estimates (e.g., Tiego et al., 2023) and improving fMRI acquisition strategies (e.g., Kundu et al., 2017; Lynch et al., 2020) may be a more fruitful strategy for addressing this question. Ongoing refinement of post-acquisition denoising procedures may offer little benefit in this regard.

### Limitations and Considerations

Our analysis centred on pre-processing steps that follow an initial minimal processing pipeline, as we anticipated that these would produce the most significant differences in FC and brain-behaviour effect sizes. Recent findings suggest that decisions made during the minimal preprocessing phase—such as the choice of registration template or processing package—can also have a substantial impact on the resulting FC (Li et al., 2024). We used the same normalization template in all our analyses, but it is possible that such choices also influence the strength of BWAS effect sizes.

Doubts have recently been raised about the efficacy of QC-FC correlations in gauging denoising efficacy, as they can increase following censoring, which specifically removes high-motion frames from the data (Williams et al., 2022). Multiple other benchmarks have been proposed (see Parkes et al., 2017 for an overview) but, here, we relied on QC-FC correlations as they are the most widely studied benchmark metric and they facilitate comparison with past work (Ciric et al., 2017; Parkes et al., 2017).

A major limiting factor on the effect sizes achievable in BWAS is the validity and reliability of the behavioural measures. Reliance on raw test or summed scale scores assumes that these measures are conceptually sound and accurate representations of the behavioural construct of interest (Tiego et al., 2023). It is well established in psychometrics that self-report measures of behaviour exhibit low reliability in capturing the target behaviour (Dang et al., 2020) and this poor reliability will attenuate brain-behaviour correlations (Saccenti et al., 2020). The observed predictive performance of ∼0.1 for most measures in this study might simply reflect reliance on the simplest possible behavioural estimates. Future research should use more advanced modelling of behavioural measure (e.g., Tiego et al., 2023) to explore whether BWAS effect sizes can be improved beyond the magnitudes observed in this study.

### Conclusions

We compared six different fMRI pre-processing procedures, aggregated into 14 distinct pipelines, across three independent datasets. Pipelines incorporating ICA-FIX with GSR generally performed well in terms of denoising efficacy and behavioural predictions, but no single pipeline was optimal for all benchmarks across all datasets, and BWAS effect sizes remained small, regardless of denoising efficacy. Our findings point to a need to identify the conditions that determine which pipeline may be optimally suited for a given dataset and to focus attention on alternative data acquisition and behavioural modelling strategies for enhancing the magnitude of brain-behaviour correlations.

## References

Aquino, K. M., Fulcher, B. D., Parkes, L., Sabaroedin, K., & Fornito, A. (2020). Identifying and removing widespread signal deflections from fMRI data: Rethinking the global signal regression problem. NeuroImage, 212, 116614. 10.1016/j.neuroimage.2020.116614

Avants, B., Tustison, N. J., & Song, G. (2009). Advanced normalization tools: V1. 0. The Insight Journal. 10.54294/uvnhin

Aref, S., & Mostajabdaveh, M. (2024). Analyzing modularity maximization in approximation, heuristic, and graph neural network algorithms for community detection. Journal of Computational Science, 78, 102283. 10.1016/j.jocs.2024.102283

Baniqued, P. L., Gallen, C. L., Kranz, M. B., Kramer, A. F., & D’Esposito, M. (2019). Brain network modularity predicts cognitive training-related gains in young adults. Neuropsychologia, 131, 205–215. 10.1016/j.neuropsychologia.2019.05.021

Beckmann, C. F., & Smith, S. M. (2004). Probabilistic independent component analysis for functional magnetic resonance imaging. IEEE Transactions on Medical Imaging, 23(2), 137–152. 10.1109/TMI.2003.822821

Betzel, R. F. (2020). Community detection in network neuroscience. 10.48550/ARXIV.2011.06723

Betzel, R. F., Medaglia, J. D., Papadopoulos, L., Baum, G. L., Gur, R., Gur, R., Roalf, D., Satterthwaite, T. D., & Bassett, D. S. (2017). The modular organization of human anatomical brain networks: Accounting for the cost of wiring. Network Neuroscience, 1(1), 42–68. 10.1162/NETN_a_00002

Birn, R. M., Diamond, J. B., Smith, M. A., & Bandettini, P. A. (2006). Separating respiratory-variation-related fluctuations from neuronal-activity-related fluctuations in fMRI. NeuroImage, 31(4), 1536–1548. 10.1016/j.neuroimage.2006.02.048

Burgess, G. C., Kandala, S., Nolan, D., Laumann, T. O., Power, J. D., Adeyemo, B., Harms, M. P., Petersen, S. E., & Barch, D. M. (2016). Evaluation of Denoising Strategies to Address Motion-Correlated Artifacts in Resting-State Functional Magnetic Resonance Imaging Data from the Human Connectome Project. Brain Connectivity, 6(9), 669–680. 10.1089/brain.2016.0435

Byrge, L., & Kennedy, D. P. (2018). Identifying and characterizing systematic temporally-lagged BOLD artifacts. NeuroImage, 171, 376–392. 10.1016/j.neuroimage.2017.12.082

Carver, C. S., & White, T. L. (1994). Behavioral inhibition, behavioral activation, and affective responses to impending reward and punishment: The BIS/BAS Scales. Journal of Personality and Social Psychology, 67(2), 319–333. 10.1037/0022-3514.67.2.319

Chen, J., Tam, A., Kebets, V., Orban, C., Ooi, L. Q. R., Asplund, C. L., Marek, S., Dosenbach, N. U. F., Eickhoff, S. B., Bzdok, D., Holmes, A. J., & Yeo, B. T. T. (2022). Shared and unique brain network features predict cognitive, personality, and mental health scores in the ABCD study. Nature Communications, 13(1), 2217. 10.1038/s41467-022-29766-8

Chyzhyk, D., Varoquaux, G., Milham, M., & Thirion, B. (2022). How to remove or control confounds in predictive models, with applications to brain biomarkers. GigaScience, 11. 10.1093/gigascience/giac014

Ciric, R., Wolf, D. H., Power, J. D., Roalf, D. R., Baum, G. L., Ruparel, K., Shinohara, R. T., Elliott, M. A., Eickhoff, S. B., Davatzikos, C., Gur, R. C., Gur, R. E., Bassett, D. S., & Satterthwaite, T. D. (2017). Benchmarking of participant-level confound regression strategies for the control of motion artifact in studies of functional connectivity. NeuroImage, 154, 174–187. 10.1016/j.neuroimage.2017.03.020

Costa Jr., P. T., & McCrae, R. R. (2000). Neo Personality Inventory. In Encyclopedia of psychology, Vol. 5. (pp. 407–409). American Psychological Association. 10.1037/10520-172

Cox, R. W. (1996). Afni: Software for analysis and visualization of functional magnetic resonance neuroimages. Computers and Biomedical Research, 29(3), 162–173. 10.1006/cbmr.1996.0014

Dang, J., King, K. M., & Inzlicht, M. (2020). Why are self-report and behavioral measures weakly correlated? Trends in Cognitive Sciences, 24(4), 267–269. 10.1016/j.tics.2020.01.007

DeYoung, C. G., Hilger, K., Hanson, J. L., Abend, R., Allen, T., Beaty, R., Blain, S. D., Chavez, R. S., Engel, S. A., Ma, F., Fornito, A., Genç, E., Goghari, V., Grazioplene, R. G., Homan, P., Joyner, K., Kaczkurkin, A. N., Latzman, R. D., Martin, E. A., … Wacker, J. (2024). Beyond Increasing Sample Sizes: Optimizing Effect Sizes in Neuroimaging Research on Individual Differences. OSF. 10.31219/osf.io/bjn62

Esteban, O., Markiewicz, C. J., Blair, R. W., Moodie, C. A., Isik, A. I., Erramuzpe, A., Kent, J. D., Goncalves, M., DuPre, E., Snyder, M., Oya, H., Ghosh, S. S., Wright, J., Durnez, J., Poldrack, R. A., & Gorgolewski, K. J. (2019). Fmriprep: A robust preprocessing pipeline for functional mri. Nature Methods, 16(1), 111–116. 10.1038/s41592-018-0235-4

Fornito, A., & Bullmore, E. T. (2010). What can spontaneous fluctuations of the blood oxygenation-level-dependent signal tell us about psychiatric disorders? Current Opinion in Psychiatry, 23(3), 239–249. 10.1097/YCO.0b013e328337d78d

Fox, M. D., Snyder, A. Z., Vincent, J. L., Corbetta, M., Van Essen, D. C., & Raichle, M. E. (2005). The human brain is intrinsically organized into dynamic, anticorrelated functional networks. Proceedings of the National Academy of Sciences, 102(27), 9673–9678. 10.1073/pnas.0504136102

Fox, M. D., Zhang, D., Snyder, A. Z., & Raichle, M. E. (2009). The global signal and observed anticorrelated resting state brain networks. Journal of Neurophysiology, 101(6), 3270–3283. 10.1152/jn.90777.2008

Gershon, R. C., Wagster, M. V., Hendrie, H. C., Fox, N. A., Cook, K. F., & Nowinski, C. J. (2013). NIH Toolbox for Assessment of Neurological and Behavioral Function. Neurology, *80*(11 Suppl 3), S2–S6. 10.1212/WNL.0b013e3182872e5f

Glasser, M. F., Sotiropoulos, S. N., Wilson, J. A., Coalson, T. S., Fischl, B., Andersson, J. L., Xu, J., Jbabdi, S., Webster, M., Polimeni, J. R., Van Essen, D. C., & Jenkinson, M. (2013). The minimal preprocessing pipelines for the Human Connectome Project. NeuroImage, 80, 105–124. 10.1016/j.neuroimage.2013.04.127

Gonzalez-Castillo, J., Handwerker, D. A., Robinson, M. E., Hoy, C. W., Buchanan, L. C., Saad, Z. S., & Bandettini, P. A. (2014). The spatial structure of resting state connectivity stability on the scale of minutes. Frontiers in Neuroscience, 8. 10.3389/fnins.2014.00138

Gordon, E. M., Laumann, T. O., Gilmore, A. W., Newbold, D. J., Greene, D. J., Berg, J. J., Ortega, M., Hoyt-Drazen, C., Gratton, C., Sun, H., Hampton, J. M., Coalson, R. S., Nguyen, A. L., McDermott, K. B., Shimony, J. S., Snyder, A. Z., Schlaggar, B. L., Petersen, S. E., Nelson, S. M., & Dosenbach, N. U. F. (2017). Precision functional mapping of individual human brains. Neuron, 95(4), 791–807.e7. 10.1016/j.neuron.2017.07.011

Gorgolewski, K. J., Durnez, J., & Poldrack, R. A. (2017). Preprocessed consortium for neuropsychiatric phenomics dataset. F1000R*esearch*, *6*, 1262. 10.12688/f1000research.11964.2

Greve, D. N., & Fischl, B. (2009). Accurate and robust brain image alignment using boundary-based registration. NeuroImage, 48(1), 63–72. 10.1016/j.neuroimage.2009.06.060

Griffanti, L., Salimi-Khorshidi, G., Beckmann, C. F., Auerbach, E. J., Douaud, G., Sexton, C. E., Zsoldos, E., Ebmeier, K. P., Filippini, N., Mackay, C. E., Moeller, S., Xu, J., Yacoub, E., Baselli, G., Ugurbil, K., Miller, K. L., & Smith, S. M. (2014). ICA-based artefact removal and accelerated fMRI acquisition for improved resting state network imaging. NeuroImage, 95, 232–247. 10.1016/j.neuroimage.2014.03.034

Gur, R. C., Richard, J., Hughett, P., Calkins, M. E., Macy, L., Bilker, W. B., Brensinger, C., & Gur, R. E. (2010). A cognitive neuroscience based computerized battery for efficient measurement of individual differences: Standardization and initial construct validation. Journal of Neuroscience Methods, 187(2), 254–262. 10.1016/j.jneumeth.2009.11.017

Hariri, A. R. (2009). The neurobiology of individual differences in complex behavioral traits. Annual Review of Neuroscience, 32(1), 225–247. 10.1146/annurev.neuro.051508.135335

He, T., Kong, R., Holmes, A. J., Nguyen, M., Sabuncu, M. R., Eickhoff, S. B., Bzdok, D., Feng, J., & Yeo, B. T. T. (2020). Deep neural networks and kernel regression achieve comparable accuracies for functional connectivity prediction of behavior and demographics. NeuroImage, 206, 116276. 10.1016/j.neuroimage.2019.116276

He, Y., Wang, J., Wang, L., Chen, Z. J., Yan, C., Yang, H., Tang, H., Zhu, C., Gong, Q., Zang, Y., & Evans, A. C. (2009). Uncovering intrinsic modular organization of spontaneous brain activity in humans. PLoS ONE, 4(4), e5226. 10.1371/journal.pone.0005226

Holmes, A. J., Hollinshead, M. O., O’Keefe, T. M., Petrov, V. I., Fariello, G. R., Wald, L. L., Fischl, B., Rosen, B. R., Mair, R. W., Roffman, J. L., Smoller, J. W., & Buckner, R. L. (2015). Brain Genomics Superstruct Project initial data release with structural, functional, and behavioral measures. Scientific Data, 2(1), 150031. 10.1038/sdata.2015.31

Jenkinson, M., Bannister, P., Brady, M., & Smith, S. (2002). Improved optimization for the robust and accurate linear registration and motion correction of brain images. NeuroImage, 17(2), 825–841. 10.1006/nimg.2002.1132

Kong, R., Li, J., Orban, C., Sabuncu, M. R., Liu, H., Schaefer, A., Sun, N., Zuo, X.-N., Holmes, A. J., Eickhoff, S. B., & Yeo, B. T. T. (2019). Spatial topography of individual-specific cortical networks predicts human cognition, personality, and emotion. Cerebral Cortex, 29(6), 2533–2551. 10.1093/cercor/bhy123

Krämer, C., Stumme, J., Da Costa Campos, L., Dellani, P., Rubbert, C., Caspers, J., Caspers, S., & Jockwitz, C. (2023). Prediction of cognitive performance differences in older age from multimodal neuroimaging data. GeroScience. 10.1007/s11357-023-00831-4

Li, X., Bianchini Esper, N., Ai, L., Giavasis, S., Jin, H., Feczko, E., Xu, T., Clucas, J., Franco, A., Sólon Heinsfeld, A., Adebimpe, A., Vogelstein, J. T., Yan, C.-G., Esteban, O., Poldrack, R. A., Craddock, C., Fair, D., Satterthwaite, T., Kiar, G., & Milham, M. P. (2024). Moving beyond processing- and analysis-related variation in resting-state functional brain imaging. Nature Human Behaviour, 1–15. 10.1038/s41562-024-01942-4

Li, J., Kong, R., Liégeois, R., Orban, C., Tan, Y., Sun, N., Holmes, A. J., Sabuncu, M. R., Ge, T., & Yeo, B. T. T. (2019). Global signal regression strengthens association between resting-state functional connectivity and behavior. NeuroImage, 196, 126–141. 10.1016/j.neuroimage.2019.04.016

Lynch, C. J., Power, J. D., Scult, M. A., Dubin, M., Gunning, F. M., & Liston, C. (2020). Rapid Precision Functional Mapping of Individuals Using Multi-Echo fMRI. Cell Reports, 33(12), 108540. 10.1016/j.celrep.2020.108540

Marek, S., Tervo-Clemmens, B., Calabro, F. J., Montez, D. F., Kay, B. P., Hatoum, A. S., Donohue, M. R., Foran, W., Miller, R. L., Hendrickson, T. J., Malone, S. M., Kandala, S., Feczko, E., Miranda-Dominguez, O., Graham, A. M., Earl, E. A., Perrone, A. J., Cordova, M., Doyle, O., … Dosenbach, N. U. F. (2022). Reproducible brain-wide association studies require thousands of individuals. Nature, 603(7902), 654–660. 10.1038/s41586-022-04492-9

Mascali, D., Moraschi, M., DiNuzzo, M., Tommasin, S., Fratini, M., Gili, T., Wise, R. G., Mangia, S., Macaluso, E., & Giove, F. (2021). Evaluation of denoising strategies for task-based functional connectivity: Equalizing residual motion artifacts between rest and cognitively demanding tasks. Human Brain Mapping, 42(6), 1805–1828. 10.1002/hbm.25332

Matsui, T., Murakami, T., & Ohki, K. (2016). Transient neuronal coactivations embedded in globally propagating waves underlie resting-state functional connectivity. Proceedings of the National Academy of Sciences, 113(23), 6556–6561. 10.1073/pnas.1521299113

Mueller, S., Wang, D., Fox, M. D., Yeo, B. T. T., Sepulcre, J., Sabuncu, M. R., Shafee, R., Lu, J., & Liu, H. (2013). Individual variability in functional connectivity architecture of the human brain. Neuron, 77(3), 586–595. 10.1016/j.neuron.2012.12.028

Newman, M. E. J. (2006). Modularity and community structure in networks. Proceedings of the National Academy of Sciences, 103(23), 8577–8582. 10.1073/pnas.0601602103

Nikolaidis, A., Chen, A. A., He, X., Shinohara, R., Vogelstein, J., Milham, M., & Shou, H. (2022). Suboptimal phenotypic reliability impedes reproducible human neuroscience (p. 2022.07.22.501193). bioRxiv. 10.1101/2022.07.22.501193

Ogawa, S., Menon, R. S., Kim, S. G., & Ugurbil, K. (1998). On the characteristics of functional magnetic resonance imaging of the brain. Annual Review of Biophysics and Biomolecular Structure, 27, 447–474. 10.1146/annurev.biophys.27.1.447

Ooi, L. Q. R., Chen, J., Zhang, S., Kong, R., Tam, A., Li, J., Dhamala, E., Zhou, J. H., Holmes, A. J., & Yeo, B. T. T. (2022). Comparison of individualized behavioral predictions across anatomical, diffusion and functional connectivity MRI. NeuroImage, 263, 119636. 10.1016/j.neuroimage.2022.119636

Ooi, L. Q. R., Orban, C., Zhang, S., Nichols, T. E., Tan, T. W. K., Kong, R., Marek, S., Dosenbach, N. U., Laumann, T., Gordon, E. M., Yap, K. H., Ji, F., Chong, J. S. X., Chen, C., An, L., Franzmeier, N., Roemer, S. N., Hu, Q., Ren, J., … Initiative, A. D. N. (2024). MRI economics: Balancing sample size and scan duration in brain wide association studies. bioRxiv, 2024.02.16.580448. 10.1101/2024.02.16.580448

Parkes, L., Fulcher, B., Yücel, M., & Fornito, A. (2017). *An evaluation of the efficacy, reliability, and sensitivity of motion correction strategies for resting-state functional MRI* [Preprint]. Neuroscience. 10.1101/156380

Poldrack, R. A., Congdon, E., Triplett, W., Gorgolewski, K. J., Karlsgodt, K. H., Mumford, J. A., Sabb, F. W., Freimer, N. B., London, E. D., Cannon, T. D., & Bilder, R. M. (2016). A phenome-wide examination of neural and cognitive function. Scientific Data, 3(1), 160110. 10.1038/sdata.2016.110

Power, J. D., Barnes, K. A., Snyder, A. Z., Schlaggar, B. L., & Petersen, S. E. (2012). Spurious but systematic correlations in functional connectivity MRI networks arise from subject motion. NeuroImage, 59(3), 2142–2154. 10.1016/j.neuroimage.2011.10.018

Power, J. D., Lynch, C. J., Silver, B. M., Dubin, M. J., Martin, A., & Jones, R. M. (2019). Distinctions among real and apparent respiratory motions in human fMRI data. NeuroImage, 201, 116041. 10.1016/j.neuroimage.2019.116041

Power, J. D., Mitra, A., Laumann, T. O., Snyder, A. Z., Schlaggar, B. L., & Petersen, S. E. (2014). Methods to detect, characterize, and remove motion artifact in resting state fMRI. NeuroImage, 84, 320–341. 10.1016/j.neuroimage.2013.08.048

Power, J. D., Plitt, M., Laumann, T. O., & Martin, A. (2017). Sources and implications of whole-brain fMRI signals in humans. NeuroImage, 146, 609–625. 10.1016/j.neuroimage.2016.09.038

Power, J. D., Schlaggar, B. L., & Petersen, S. E. (2015). Recent progress and outstanding issues in motion correction in resting state fMRI. NeuroImage, 105, 536–551. 10.1016/j.neuroimage.2014.10.044

Pruim, R. H. R., Mennes, M., Van Rooij, D., Llera, A., Buitelaar, J. K., & Beckmann, C. F. (2015). ICA-AROMA: A robust ICA-based strategy for removing motion artifacts from fMRI data. NeuroImage, 112, 267–277. 10.1016/j.neuroimage.2015.02.064

Rubinov, M., & Sporns, O. (2010). Complex network measures of brain connectivity: Uses and interpretations. NeuroImage, 52(3), 1059–1069. 10.1016/j.neuroimage.2009.10.003

Saad, Z. S., Gotts, S. J., Murphy, K., Chen, G., Jo, H. J., Martin, A., & Cox, R. W. (2012). Trouble at rest: How correlation patterns and group differences become distorted after global signal regression. Brain Connectivity, 2(1), 25–32. 10.1089/brain.2012.0080

Saccenti, E., Hendriks, M. H. W. B., & Smilde, A. K. (2020). Corruption of the Pearson correlation coefficient by measurement error and its estimation, bias, and correction under different error models. Scientific Reports, 10(1), 438. 10.1038/s41598-019-57247-4

Salimi-Khorshidi, G., Douaud, G., Beckmann, C. F., Glasser, M. F., Griffanti, L., & Smith, S. M. (2014). Automatic denoising of functional MRI data: Combining independent component analysis and hierarchical fusion of classifiers. NeuroImage, 90, 449–468. 10.1016/j.neuroimage.2013.11.046

Satterthwaite, T. D., Elliott, M. A., Gerraty, R. T., Ruparel, K., Loughead, J., Calkins, M. E., Eickhoff, S. B., Hakonarson, H., Gur, R. C., Gur, R. E., & Wolf, D. H. (2013). An improved framework for confound regression and filtering for control of motion artifact in the preprocessing of resting-state functional connectivity data. NeuroImage, 64, 240–256. 10.1016/j.neuroimage.2012.08.052

Scheel, N., Keller, J. N., Binder, E. F., Vidoni, E. D., Burns, J. M., Thomas, B. P., Stowe, A. M., Hynan, L. S., Kerwin, D. R., Vongpatanasin, W., Rossetti, H., Cullum, C. M., Zhang, R., & Zhu, D. C. (2022). Evaluation of noise regression techniques in resting-state fMRI studies using data of 434 older adults. Frontiers in Neuroscience, 16, 1006056. 10.3389/fnins.2022.1006056

Schölvinck, M. L., Maier, A., Ye, F. Q., Duyn, J. H., & Leopold, D. A. (2010). Neural basis of global resting-state fMRI activity. Proceedings of the National Academy of Sciences, 107(22), 10238–10243. 10.1073/pnas.0913110107

Shmueli, K., Van Gelderen, P., De Zwart, J. A., Horovitz, S. G., Fukunaga, M., Jansma, J. M., & Duyn, J. H. (2007). Low-frequency fluctuations in the cardiac rate as a source of variance in the resting-state fMRI BOLD signal. NeuroImage, 38(2), 306–320. 10.1016/j.neuroimage.2007.07.037

Siegel, J. S., Mitra, A., Laumann, T. O., Seitzman, B. A., Raichle, M., Corbetta, M., & Snyder, A. Z. (2017). Data quality influences observed links between functional connectivity and behavior. Cerebral Cortex, 27(9), 4492–4502. 10.1093/cercor/bhw253

Siegel, J. S., Power, J. D., Dubis, J. W., Vogel, A. C., Church, J. A., Schlaggar, B. L., & Petersen, S. E. (2014). Statistical improvements in functional magnetic resonance imaging analyses produced by censoring high-motion data points. Human Brain Mapping, 35(5), 1981–1996. 10.1002/hbm.22307

Smith, S. M. (2002). Fast robust automated brain extraction. Human Brain Mapping, 17(3), 143–155. 10.1002/hbm.10062

Smith, S. M., Fox, P. T., Miller, K. L., Glahn, D. C., Fox, P. M., Mackay, C. E., Filippini, N., Watkins, K. E., Toro, R., Laird, A. R., & Beckmann, C. F. (2009). Correspondence of the brain’s functional architecture during activation and rest. Proceedings of the National Academy of Sciences, 106(31), 13040–13045. 10.1073/pnas.0905267106

Smith, S. M., Jenkinson, M., Woolrich, M. W., Beckmann, C. F., Behrens, T. E. J., Johansen-Berg, H., Bannister, P. R., De Luca, M., Drobnjak, I., Flitney, D. E., Niazy, R. K., Saunders, J., Vickers, J., Zhang, Y., De Stefano, N., Brady, J. M., & Matthews, P. M. (2004). Advances in functional and structural MR image analysis and implementation as FSL. NeuroImage, 23 *Suppl 1*, S208–219. 10.1016/j.neuroimage.2004.07.051

Spielberger, C. D., Gorsuch, R. L., & Lushene. R. E. (1970). Manual for the State-Trait Anxiety Inventory (Self-evaluation Questionnaire). Consulting Psychologists Press.

Sporns, O., & Betzel, R. F. (2016). Modular Brain Networks. Annual Review of Psychology, 67, 613–640. 10.1146/annurev-psych-122414-033634

Teeuw, J., Hulshoff Pol, H. E., Boomsma, D. I., & Brouwer, R. M. (2021). Reliability modelling of resting-state functional connectivity. NeuroImage, 231, 117842. 10.1016/j.neuroimage.2021.117842

Tiego, J., Martin, E. A., DeYoung, C. G., Hagan, K., Cooper, S. E., Pasion, R., Satchell, L., Shackman, A. J., Bellgrove, M. A., Fornito, A., Abend, R., Goulter, N., Eaton, N. R., Kaczkurkin, A. N., & Nusslock, R. (2023). Precision behavioral phenotyping as a strategy for uncovering the biological correlates of psychopathology. Nature Mental Health, 1(5), 304–315. 10.1038/s44220-023-00057-5

Tustison, N. J., Avants, B. B., Cook, P. A., Egan, A., Yushkevich, P. A., & Gee, J. C. (2010). N4itk: Improved n3 bias correction. IEEE Transactions on Medical Imaging, 29(6), 1310– 1320. 10.1109/TMI.2010.2046908

Van Dijk, K. R. A., Sabuncu, M. R., & Buckner, R. L. (2012). The influence of head motion on intrinsic functional connectivity MRI. NeuroImage, 59(1), 431–438. 10.1016/j.neuroimage.2011.07.044

Van Essen, D. C., Smith, S. M., Barch, D. M., Behrens, T. E. J., Yacoub, E., & Ugurbil, K. (2013). The wu-minn human connectome project: An overview. NeuroImage, 80, 62–79. 10.1016/j.neuroimage.2013.05.041

Van Essen, D. C., Ugurbil, K., Auerbach, E., Barch, D., Behrens, T. E. J., Bucholz, R., Chang, A., Chen, L., Corbetta, M., Curtiss, S. W., Della Penna, S., Feinberg, D., Glasser, M. F., Harel, N., Heath, A. C., Larson-Prior, L., Marcus, D., Michalareas, G., Moeller, S., … Yacoub, E. (2012). The human connectome project: A data acquisition perspective. NeuroImage, 62(4), 2222–2231. 10.1016/j.neuroimage.2012.02.018

Weinberger, A. B., Cortes, R. A., Betzel, R. F., & Green, A. E. (2022). *Exploring functional brain network modularity in educational contexts* [Preprint]. Neuroscience. 10.1101/2022.01.06.475275

Williams, J. C., Tubiolo, P. N., Luceno, J. R., & Van Snellenberg, J. X. (2022). Advancing motion denoising of multiband resting-state functional connectivity fMRI data. NeuroImage, 249, 118907. 10.1016/j.neuroimage.2022.118907

Xifra-Porxas, A., Kassinopoulos, M., Prokopiou, P., Boudrias, M.-H., & Mitsis, G. D. (2024). Does global signal regression alter fMRI connectivity patterns related to EEG activity? An EEG-fMRI study in humans (p. 2024.04.18.590163). bioRxiv. 10.1101/2024.04.18.590163

