## Supplementary Information for "The efficacy of resting-state fMRI denoising pipelines for motion correction and behavioural prediction"

### Supplementary Methods

*1. Modularity Maximisation*

In the main text, we evaluated the performance of diffuse cluster estimation and regression (DiCER) using between 1 and 5 iterations. However, the same number of iterations is applied to all individuals, which may limit the performance of this algorithm. For instance, few iterations may be too lenient in individuals with high-noise data, whereas multiple iterations may be too aggressive in individuals with low-noise data (Aquino et al., 2020).

We developed a simply heuristic for tailoring the number of DiCER iterations to an individual dataset that is based on maximizing the modularity of the resulting functional coupling (FC) network. Modularity refers to the degree to which a network can be divided into cohesive communities, such that networks with high modularity have stronger FC within communities relative to a null model (Newman, 2006). In datasets with many wide-spread signal deflections (WSDs), FC networks are globally coherent and the network shows low modularity; similarly, if denoising is too aggressive, the network will show little structure, which also reduces network modularity. This implies that there may be an optimal number of DiCER iterations interposed between these two extremes that is associated with some maximal degree of modularity.

We used the Louvain algorithm (Blondel et al., 2008), as implemented in the Brain Connectivity Toolbox adapted for python v0.61 (Rubinov and Sporns, 2010) to find a partition of each person's FC network that seeks to maximize the $Q$-statistic, which quantifies the difference between intra-module FC and chance expectations,

$$Q = \frac{1}{2\mu}\sum_{i,j}\left(A_{ij} - \gamma V_{ij}\right)\delta\left(m_{i,}m_j\right), \tag{1}$$

where $A_{ij}$ refers to the FC weight between nodes $i$ and $j$, $\mu$ is the total weight of the network, $V_{ij} = \frac{k_i k_j}{2\mu}$ is the FC weight of the corresponding null model, with $k_i$ and $k_j$ corresponding to the weighted degree of nodes $i$ and $j$,. The Kronecker delta function $\delta(m_i, m_j) = 1$ if nodes $i$ and $j$ are in the same module and zero otherwise. The resolution parameter, $\gamma$, tunes the size of the resulting modules such that larger $\gamma$ yields smaller modules.

The modularity maximization algorithm was run 100 times on individual FC matrices. The resulting 100 node assignments were then subjected to a consensus clustering (Lancichinetti and Fortunato, 2012) and the final $Q$ value for a given FC matrix was computed from the consensus partition. This algorithm was executed across every iteration of DiCER, and the maximum $Q$ value across iterations was chosen as the optimal number of DiCER iterations.

### 2. *Pipelines*

*Supplementary Table 1.* Different pre-processing pipelines employed in the our analysis.

| *Pipeline Number* | *Pipeline steps* |
|---|---|
| *1.* | 24P |
| *2.* | 28P |
| *3.* | AROMA + 28P |
| *4.* | AROMA + 28P + Censoring |
| *5.* | FIX + 28P |
| *6.* | FIX + 28P + Censoring |
| *7.* | AROMA + 28P + GSR |
| *8.* | AROMA + 28P + GSR + Censoring |
| *9.* | FIX + 28P + GSR |
| *10.* | FIX + 28P + GSR + Censoring |
| *11.* | AROMA + 28P + DiCER |
| *12.* | AROMA + 28P + DiCER + Censoring |
| *13.* | FIX + 28P + DiCER |
| *14.* | FIX + 28P + DiCER + Censoring |

### *3. Behavioural Variables*

*Supplementary Table 2.* Behavioural variables used in the Kernel Ridge Regression (KRR) for the GSP dataset. For each variable, the corresponding field names in the GSP behavioural csv file are shown together with whether they belonged to the cognition or personality domain. The table has been adapted from Li et al. (2019).

| *Description* | *Name in the GSP dataset* | *Cognitive* | *Personality* | *Others* |
|---|---|---|---|---|
| *Flanker Accuracy* | Flanker_S_CORRpc | ✓ | | |
| *Mental Rotation Accuracy* | Average of MenRot_80_CORRpc, MenRot_120_CORRpc, and MentRot_160_CORRpc | ✓ | | |
| *Shipley Vocabulary* | Shipley_Vocab_Raw | ✓ | | |
| *WAIS Matrix Reasoning* | Matrix_WAIS | ✓ | | |
| *Speilberger Trait Anxiety* | STAI_tAnxiety | | | ✓ |
| *Speilberger State Anxiety* | STAI_sAnxiety | | | ✓ |
| *Total Mood Disturbance* | POMS_TotMdDisturb | | | ✓ |
| *Barratt Impulsivity* | Barratt_tot | | | ✓ |
| *Mind Wandering Frequency* | MindWandering_Freq | | | ✓ |
| *NEO Neuroticism* | NEO_N | | ✓ | |
| *NEO Extraversion* | NEO_E | | ✓ | |
| *NEO Openness* | NEO_O | | ✓ | |
| *NEO Agreeableness* | NEO_A | | ✓ | |
| *NEO Conscientiousness* | NEO_C | | ✓ | |
| *Novelty Seeking* | TCI_Novelty | | ✓ | |
| *Reward Dependence* | TCI_RewardDependance | | | |
| *Harm Avoidance* | TCI_HarmAvoidance | | | ✓ |
| *Risk Taking* | DOSPERT_taking | | ✓ | |
| *Behavioural Activation Drive* | BISBAS_BAS_Drive | | | ✓ |
| *Behavioural Activation Fun* | BISBAS_BAS_Fun | | | ✓ |
| *Behavioural Activation Reward* | BISBAS_BAS_Reward | | | ✓ |
| *Behavioural Inhibition* | BISBAS_BIS | | | ✓ |

*Supplementary Table 3.* Behavioural variables used in the KRR for the HCP dataset. For each variable, the corresponding field names in the HCP behavioural csv file are shown together with whether they belonged to the cognition or personality domain. The table has been adapted from Li et al. (2019).

| *Description* | *Name in the HCP dataset* | *Cognitive* | *Personality* | *Others* |
|---|---|---|---|---|
| *Visual Episodic Memory* | PicSeq_Unadj_Score | ✓ | | |
| *Cognitive Flexibility (DCCS)* | CardSort_Unadj | ✓ | | |
| *Inhibition (Flanker Task)* | Flanker_Unadj | ✓ | | |
| *Fluid Intelligence (PMAT)* | PMAT24_A_CR | ✓ | | |
| *Vocabulary (Pronunciation)* | ReadEng_Unadj | ✓ | | |
| *Vocabulary (Picture Matching)* | PicVocab_Unadj | ✓ | | |
| *Processing Speed* | ProcSpeed_Unadj | ✓ | | |
| *Delay Discounting* | DDisc_AUC_40K | ✓ | | |
| *Spatial Orientation* | VSPLOT_TC | ✓ | | |
| *Sustained Attention – Sens.* | SCPT_SEN | ✓ | | |
| *Sustained Attention – Spec.* | SCPT_SPEC | ✓ | | |
| *Verbal Episodic Memory* | IWRD_TOT | ✓ | | |
| *Working Memory (List Sorting)* | ListSort_Unadj | ✓ | | |
| *Cognitive Status (MMSE)* | MMSE | ✓ | | |

| | | | | |
|---|---|---|---|---|
| *Sleep Quality (PSQI)* | PSQI_Score | | | ✓ |
| *Walking Endurance* | Endurance_Unadj | | | ✓ |
| *Walking Speed* | GaitSpeed_Comp | | | ✓ |
| *Manual Dexterity* | Dexterity_Unadj | | | ✓ |
| *Grip Strength* | Strength_Unadj | | | ✓ |
| *Odour Identification* | Odor_Unadj | | | ✓ |
| *Pain Interference Survey* | PainInterf_Tscore | | | ✓ |
| *Taste Intensity* | Taste_Unadj | | | ✓ |
| *Contrast Sensitivity* | Mars_Final | | | ✓ |
| *Emotional Face Matching* | Emotion_Task_Face_Acc | | | ✓ |
| *Arithmetic* | Language_Task_Math_Avg_Difficulty_Level | ✓ | | |
| *Story Comprehension* | Language_Task_Story_Avg_Difficulty_Level | ✓ | | |
| *Relational Processing* | Relational_Task_Acc | | | ✓ |
| *Social Cognition – Random* | Social_Task_Perc_Random | | | ✓ |
| *Social Cognition – Interaction* | Social_Task_Perc_TOM | | | ✓ |
| *Working Memory (N-back)* | WM_Task_Acc | ✓ | | |
| *Agreeableness (NEO)* | NEOFAC_A | | ✓ | |
| *Openness (NEO)* | NEOFAC_O | | ✓ | |
| *Conscientiousness (NEO)* | NEOFAC_C | | ✓ | |
| *Neuroticism (NEO)* | NEOFAC_N | | ✓ | |

| | | | | |
|---|---|---|---|---|
| *Extraversion (NEO)* | NEOFAC_E | | ✓ | |
| *Emotion Recognition – Total* | ER40_CR | | | ✓ |
| *Emotion Recognition – Angry* | ER40ANG | | | ✓ |
| *Emotion Recognition – Fear* | ER40FEAR | | | ✓ |
| *Emotion Recognition – Happy* | ER40HAP | | | ✓ |
| *Emotion Recognition - Neutral* | ER40NOE | | | ✓ |
| *Emotion Recognition – Sad* | ER40SAD | | | ✓ |
| *Anger – Affect* | AngAffect_Unadj | | ✓ | |
| *Anger – Hostility* | AngHostil_Unadj | | ✓ | |
| *Anger – Aggression* | AngAggr_Unadj | | ✓ | |
| *Fear – Affect* | FearAffect_Unadj | | ✓ | |
| *Fear – Somatic Arousal* | FearSomat_Unadj | | ✓ | |
| *Sadness* | Sadness_Unadj | | ✓ | |
| *Life Satisfaction* | LifeSatisf_Unadj | | | ✓ |
| *Meaning & Purpose* | MeanPurp_Unadj | | | ✓ |
| *Positive Affect* | PosAffect_Unadj | | ✓ | |
| *Friendship* | Friendship_Unadj | | | ✓ |
| *Loneliness* | Loneliness_Unadj | | | ✓ |
| *Perceived Hostility* | PercHostil_Unadj | | | ✓ |
| *Perceived Rejection* | PercReject_Unadj | | | ✓ |
| *Emotional Support* | EmotSupp_Unadj | | | ✓ |

| | | | | |
|---|---|---|---|---|
| *Instrument Support* | InstruSupp_Unadj | | | ✓ |
| *Perceived Stress* | PercStress_Unadj | | | ✓ |
| *Self-Efficacy* | SelfEff_Unadj | | | ✓ |

### Supplementary Figures

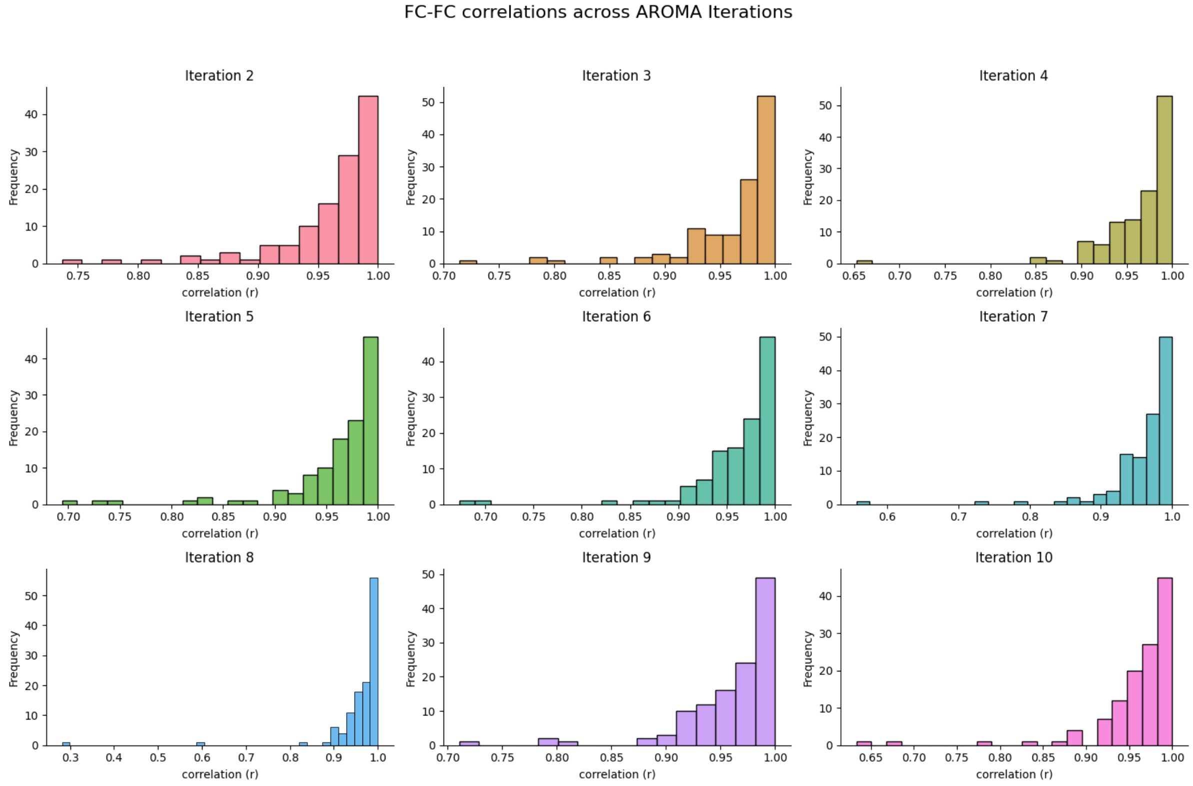


*Supplementary Figure 1.* Histograms illustrating the variability of ICA-AROMA across iterations in the CNP dataset. Each subject underwent a single ICA-AROMA iteration, where the upper triangle of their FC matrix was correlated against nine additional iterations using Pearson’s R. The plot displays the nine FC-FC correlations for each subject, revealing the distribution of individual FC changes across different runs of ICA-AROMA.

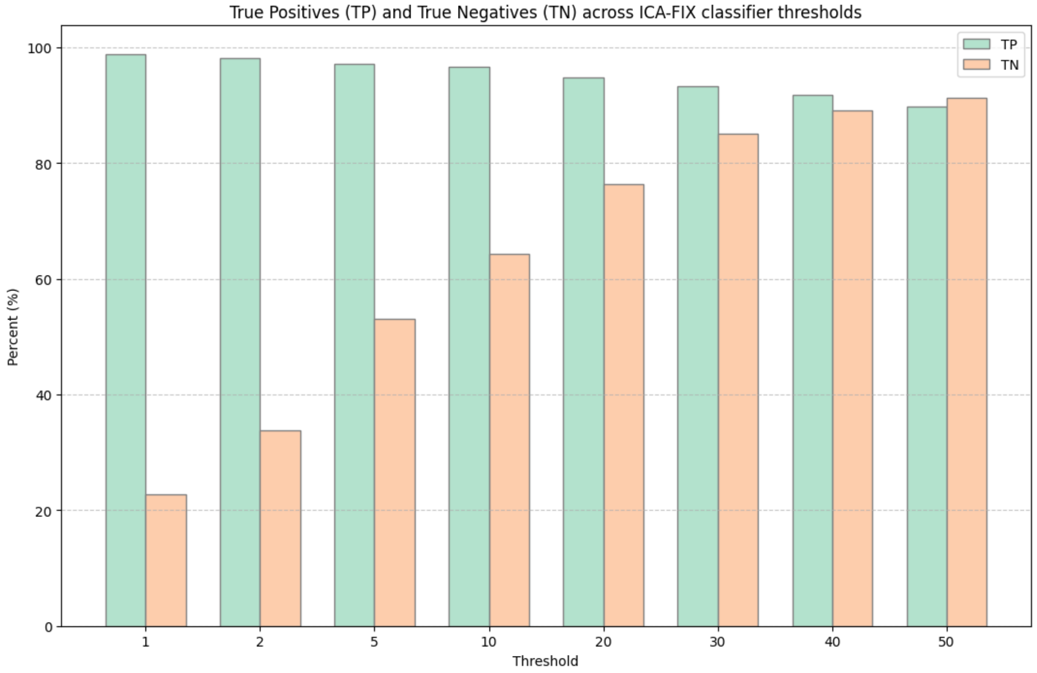


*Supplementary Figure 2.* Performance of the ICA-FIX classifier across various thresholds in the GSP dataset. Leave-one-out cross-validation was conducted on all training subjects, and the True Positive (TP) and True Negative (TN) rates were averaged across all participants for each classifier threshold. The final threshold for the test data was determined to be 40, as it optimized the rate of TN while simultaneously maintaining a high rate of TP.

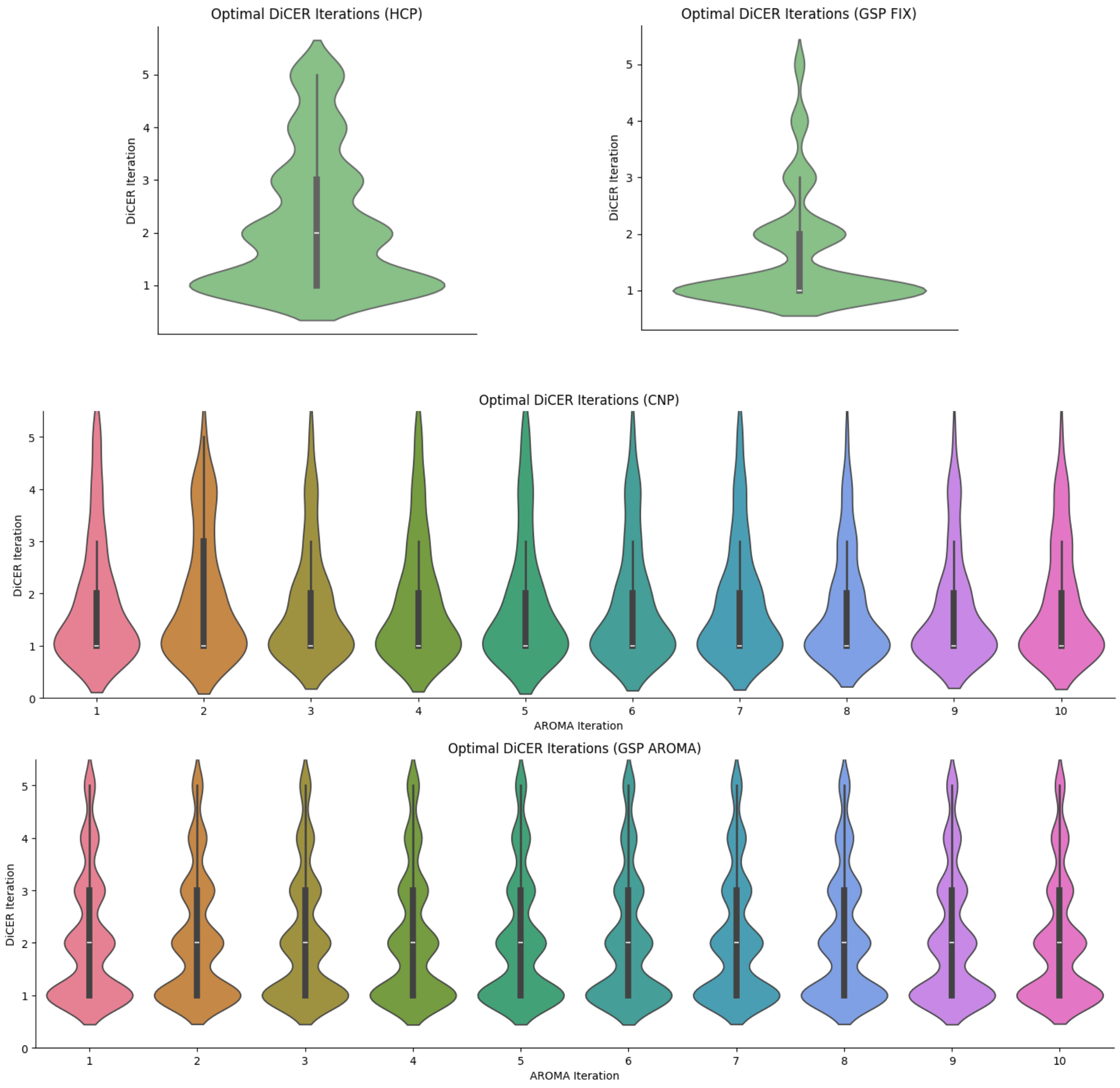


*Supplementary Figure 3.* Violin plots showing the number of iterations required by DiCER to achieve maximum modularity in each dataset. For pipelines utilizing ICA-AROMA, the maximum modularity was calculated separately for each run.

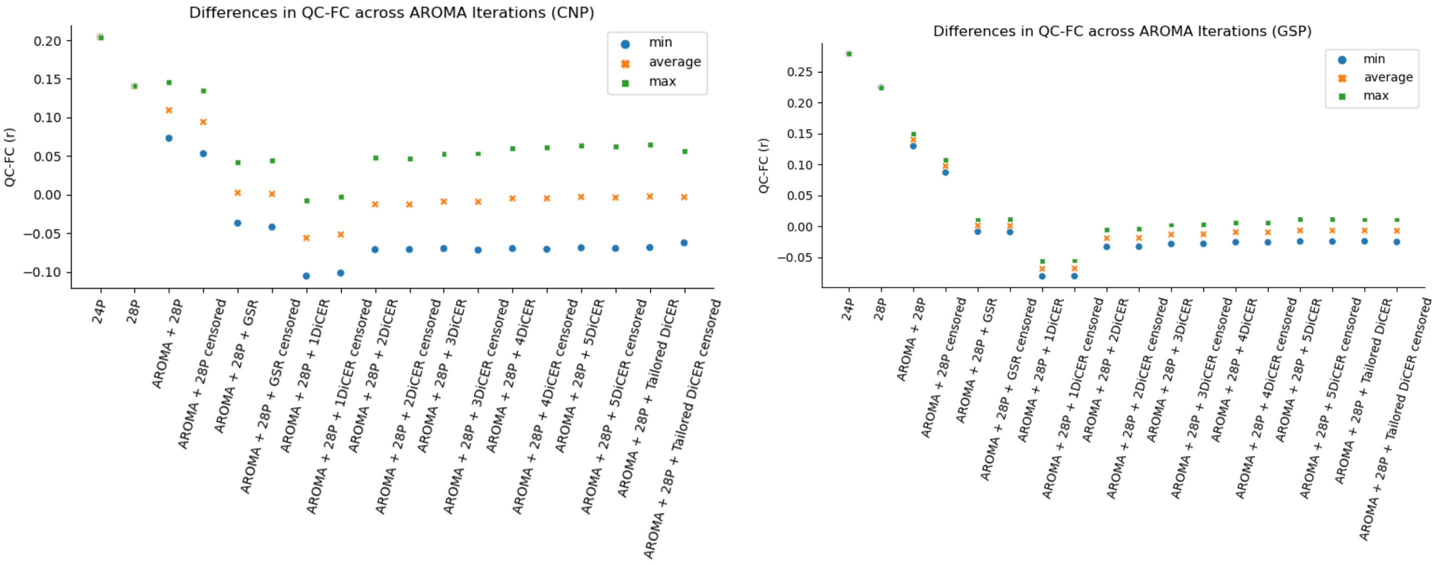


*Supplementary Figure 4.* Variability in QC-FC correlations across ICA-AROMA iterations. Scatter points depict the mean QC-FC value for each ICA-AROMA iteration, with distinctions for the lowest (blue dots), highest (green dots), and most average (orange stars) QC-FC results. The figure illustrates that differences observed across pipelines increase as more regressors are incorporated post ICA-AROMA.

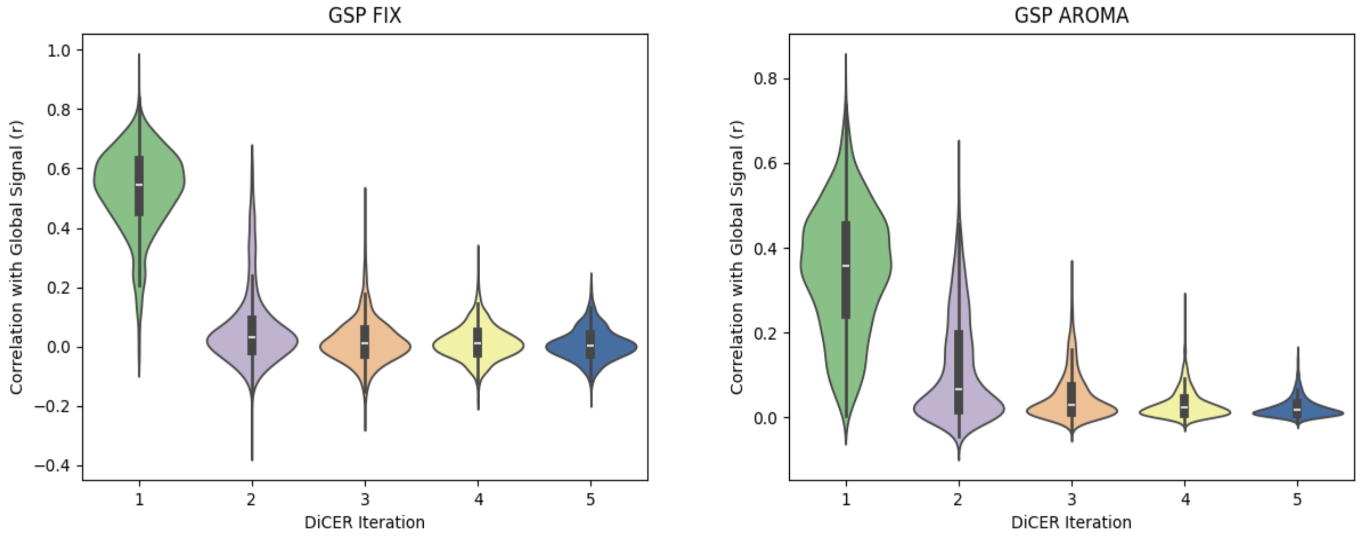


*Supplementary Figure 5.* Pearson's correlation of DiCER regressors with global signal in the GSP dataset. Violin plots depict the correlation between each subject's global signal and DiCER regressors. Shaded areas represent the kernel density estimate of each correlations distribution, while the boxes represent the median (white lines) and interquartile ranges. For pipelines incorporating ICA-AROMA, correlations were averaged across 5 iterations. The first DiCER regressor exhibits a moderate correlation with the global signal, the second displays a low correlation, while the subsequent regressors show no discernible correlation.

### References

Aquino, K. M., Fulcher, B. D., Parkes, L., Sabaroedin, K., & Fornito, A. (2020). Identifying and removing widespread signal deflections from fMRI data: Rethinking the global signal regression problem. NeuroImage, 212, 116614. https://doi.org/10.1016/j.neuroimage.2020.116614

Blondel, V. D., Guillaume, J.-L., Lambiotte, R., & Lefebvre, E. (2008). Fast unfolding of communities in large networks. *Journal of Statistical Mechanics: Theory and Experiment*, *2008*(10), P10008. https://doi.org/10.1088/1742-5468/2008/10/P10008

Lancichinetti, A., & Fortunato, S. (2012). Consensus clustering in complex networks. *Scientific Reports*, *2*(1), 336. https://doi.org/10.1038/srep00336

Newman, M. E. J. (2006). Modularity and community structure in networks. *Proceedings of the National Academy of Sciences*, *103*(23), 8577–8582. https://doi.org/10.1073/pnas.0601602103

Rubinov, M., & Sporns, O. (2010). Complex network measures of brain connectivity: Uses and interpretations. *NeuroImage*, *52*(3), 1059–1069. https://doi.org/10.1016/j.neuroimage.2009.10.003
